# Membrane-penetrating peptide from the translocation region of *Bordetella* Adenylate Cyclase Toxin prevents toxin cytotoxicity on target cells

**DOI:** 10.1101/2023.04.18.537300

**Authors:** Jone Amuategi, Rocío Alonso, Igor de la Arada, Helena Ostolaza

**Affiliations:** Department of Biochemistry and Molecular Biology, University of Basque Country (UPV/EHU); Biofisika Institute (UPV/EHU, CSIC) and Aptdo. 644, 48080 Bilbao, Spain; Fundación Biofísica Bizkaia (FBB), Barrio Sarriena s/n, 48940 Leioa, Spain

**Keywords:** Cholesterol, CRAC motif, pore-forming toxins, RTX toxin family, lipid-protein interactions

## Abstract

Adenylate cyclase toxin (ACT) is one of the main virulence factors of *Bordetella pertussis*, with crucial role in colonization of human respiratory tract. ACT toxicity on target phagocytes results from translocation of its adenylate cyclase domain and production of high cAMP levels and from pore formation. Recently, we unveiled in ACT four cholesterol-recognition motifs involved in specific interaction with membrane cholesterol, which might stabilize membrane topology of critical helices for ACT activity. Here we explore an amphipathic peptide corresponding to ACT residues 454 to 487 containing one of such CRAC motifs. We show that P454-487 penetrates into DOPC vesicles as a long and tilted α-helix, while in cholesterol presence experiments conformational changes that critically depend on the CRAC Phe-485 residue. Moreover, P454-487 is capable of blocking ACT toxicity on cells by outcompeting with the full-length toxin for membrane binding. We anticipate P454-487 may have potential clinical applicability in controlling *Bordetella* infection.

## INTRODUCTION

Whooping cough is a severe respiratory infectious disease caused by the bacterium *Bordetella pertussis* [1]. In the pre-vaccination era, whooping cough epidemics occurred extensively and presented a cyclical pattern, with incidence peaks every 2 to 5 years [2, 3]. The disease mainly affected children, representing one of the largest causes of infant mortality globally. The introduction worldwide of the vaccine in the forties led to a large decrease in the number of cases [2, 4]. However, epidemic peaks have continued being recorded every 4 to 5 years, leading the WHO to declare whooping cough re-emerging disease [4-6]. Consequently, whooping cough represents today one of the main causes of mortality due to vaccine-preventable diseases for the child population. The adenylate cyclase toxin (ACT or CyaA) is one of the important virulence factors secreted by *B. pertussis*, playing an important role in the early stages of respiratory tract colonization by this organism [7].

ACT is a multi-domain 1706-residue-long protein toxin of the RTX (**R**epeats in **T**o**X**in) family [7-10]. The first 364 amino-proximal residues conform the catalytic adenylate cyclase (AC) domain with ATP-cyclizing activity [7], whereas the C-terminal 1306 residues constitute the “RTX hemolysin” moiety endowed with an intrinsic pore-forming lytic activity [7-10]. Joining both domains is the so-called “translocation region” (TR) (amino acids ≈365-500) that is essential for the transport of the AC domain into the cytosol of target cells [11, 12]. In the RTX moiety in turn, several regions can be further delineated. The hydrophobic region (HR) (residues ≈500 to 710) contains several hydrophobic or amphipathic segments that are predicted to adopt α-helical structures and to insert into membranes to form cation-selective pores [13, 14]. The acylation region (residues 711–1005) contains two post-translational acylation sites, at lysines K860 and K983, to which two fatty acids are covalently linked by a dedicated acyltransferase (CyaC) [15, 16]. The Repeats domain (residues 1006-1706) consists of ∼40 copies of a glycine and aspartate-rich nonapeptide repeat characteristic of the RTX cytolysins [7, 8,17] and is involved in calcium binding by ACT, which is necessary for the toxin biological activity [17, 18]. The C-terminal moiety also contains the binding site for the CD11b/CD18 integrin, specific cellular receptor [19, 20], and a secretion signal located at the C-terminal extremity recognized by the dedicated Type 1 Secretion System (T1SS), made of CyaB, CyaD, and CyaE proteins [21,22].

Myeloid phagocytes that express the CD11b/CD18 integrin are believed to be the natural targets for ACT [19, 20]. However, the toxin can also bind at detectable levels to a variety of eukaryotic cell types from diverse mammalian species [23-25] and even to protein-free lipid bilayers of pure phospholipids [26]. Intoxication of target cells occurs upon high affinity binding of the RTX domain to the CD11b/CD18 integrin at the cell membrane [19]. Then, ACT inserts several helical elements of the translocation and hydrophobic regions into the lipid bilayer that subsequently lead to the translocation of the N-terminal AC domain into the target cytosol. The exact molecular mechanism followed by the AC domain to be delivered into the target cytosol is not understood and is a topic highly debated in the field. Once inside the cell the AC domain binds calmodulin and converts ATP into cAMP with a high catalytic turnover [27]. Accumulation of cAMP and depletion of ATP subvert the phagocytic functions of the immune cells leading to decrease in host defence [28]. At higher concentrations, ACT causes the permeabilization of the cellular membrane by forming oligomeric lytic pores that lead ultimately to cell death [29-31].

Binding of toxin monomers to the membrane, and their correct insertion and oligomerization into the lipid bilayer, are essential steps common in the action mechanisms of protein toxins acting at the cell membrane. Membrane lipids such as sphingomyelin and cholesterol have been shown for several protein toxins to have an essential role by favoring one or more of these steps [32-34]. In recent years, experimental evidences have shown that several RTX toxins including the leukotoxin (LtxA) from *Aggregatibacter actinomytemcomitans*, the *Escherichia coli* hemolysin (HlyA) and RtxA cytolysin from *Kingella kingae*, require membrane cholesterol for their correct lytic functions [35-37]. In a recent study from our laboratory, we showed that ACT binding to membrane cholesterol is essential for both lytic and AC translocation activities [38]. Moreover, we revealed that four functional Cholesterol Recognition Amino Acid Consensus (CRAC/CARC) motifs present in ACT primary structure are involved in regulating ACT binding to this sterol [38]. Two of the identified motifs, CARC-415 and CRAC-485, are localized in the TR which is predicted to be of helical structure, and that has been reported to be directly involved in AC domain translocation [12, 39, 40]. The other two, CARC-521 and CRAC-532, are located in two of the five predicted amphipathic/hydrophobic α-helixes of the HD that supposedly insert into the lipid bilayer and form part of the ACT pore structure [41-46].

CRAC is a linear motif with a simple consensus L/V-(X)1-5-Y/F-(X)1-5-R/K (in the N-terminus to the C-terminus direction), where (X) 1-5 represents between one and five residues of any amino acid [47, 48]. The exact reverse sequence, R/K-(X)1-5-Y/F-(X)1-5 -L/V, with a flexibility for central aromatic amino acid, which could be either Y or F, constitutes the CARC motif [48, 49]. These cholesterol-binding motifs were primarily found to be associated with transmembrane helices of mammalian integral membrane proteins and facilitate strong interaction between those protein segments and plasma membrane cholesterol [50, 51]. More recently, it has evidenced that these motifs are used also by many pathogens to recognize cholesterol. For example, they were identified in bacterial toxins such as the *A. actinomycetemcomitans* cytolethal distending toxin subunit C (CdtC) [52], and several viral proteins involved in membrane fusion, including the HIV transmembrane protein gp41, the influenza virus M2 protein and the SARS-CoV-2 spike S homotrimeric glycoprotein [53-55]. Interestingly, several CRAC motifs have been identified in the sequences of the aforementioned RTX toxins LtxA, HlyA and RtxA, reported to require cholesterol for their lytic activity [35-37], suggesting that interaction with the sterol may be conserved in this toxin family. Binding to cholesterol is often an essential initial step in the mechanism of these pathogens, as the toxin or virus transits from the aqueous extracellular environment to the hydrophobic membrane environment. Thus, disruption of this recognition process represents a possible method of inhibiting bacterial and viral pathogenesis.

Here we investigated a synthetic peptide, P454-487, that corresponds to residues 454-487 of ACT primary structure. It contains at the C-terminus the CRAC-485 motif that is involved in cholesterol recognition by the full-length toxin [38]. The aim of the study was to explore the membrane-interacting characteristics of P454-487, in particular whether is capable of binding to cholesterol-containing membranes and, consequently be able to block ACT binding and toxicity on cells. To that end, we have combined conventional FT-IR spectroscopy, two-dimensional correlation IR spectroscopy (2D-COS-IR), and ATR-IR, to analyze the conformation and orientation adopted by this peptide upon reconstitution into lipid bilayers. We show that P454-487 adopts α-helical structure in both pure DOPC and DOPC:Chol vesicles with low cholesterol concentrations, penetrating the lipid bilayer in a tilted orientation. However, in cholesterol-rich membranes the P454-487 helix experiments conformational changes that depend on the presence of an intact C-terminal CRAC motif and central Phe-485 residue. Besides, we show that P454-487 is not itself lytic, but its binding to the plasma membrane of target cells inhibits both pore formation and AC domain translocation, thus indicating that it may have a potential clinical applicability in controlling infection of *Bordetella*.

## RESULTS

### Membrane association of the P454-487 peptide to neutral lipid bilayers

We evaluated first whether the synthetic peptide P454-487 was capable of interacting with lipid membranes, and if so, whether cholesterol had any effect on the membrane association. To that end, and using a microfiltration methodology, we quantified the binding of the P454-487 peptide to LUVs made of pure DOPC (1,2-Dioleoyl-*sn*-glycero-3-phosphocholine) or of mixtures of DOPC and cholesterol (1:1 molar ratio). The amino acid sequence and characteristics of the P454-487 peptide are listed under “Methods” section.

As illustrated in Fig 1, binding of P454-487 to pure DOPC vesicles was very efficient with almost 100% of the peptide associated to the neutral LUVs. Incorporation of 50 % cholesterol in the vesicles (DOPC:Chol 1:1 molar ratio) did not cause significant variation in the association efficiency of the peptide, achieving almost 100% binding in both cases. In view of our previous data [38], we would expect a greater binding efficiency for the peptide in the cholesterol-containing vesicles due to the presence of the CRAC^485^ motif at the C-terminal end of P454-487. However, due to methodological constraints we used high peptide:lipid ratios in these experiments, and so for all the compositions the vesicles might be saturated with peptide molecules this way explaining the lack of differences in the binding extents. Taken together, the data indicate that the P454-487 peptide is able to efficiently partition into neutral lipid bilayers, both of DOPC and DOPC:Chol equimolar mixture.

**Figure 1.**
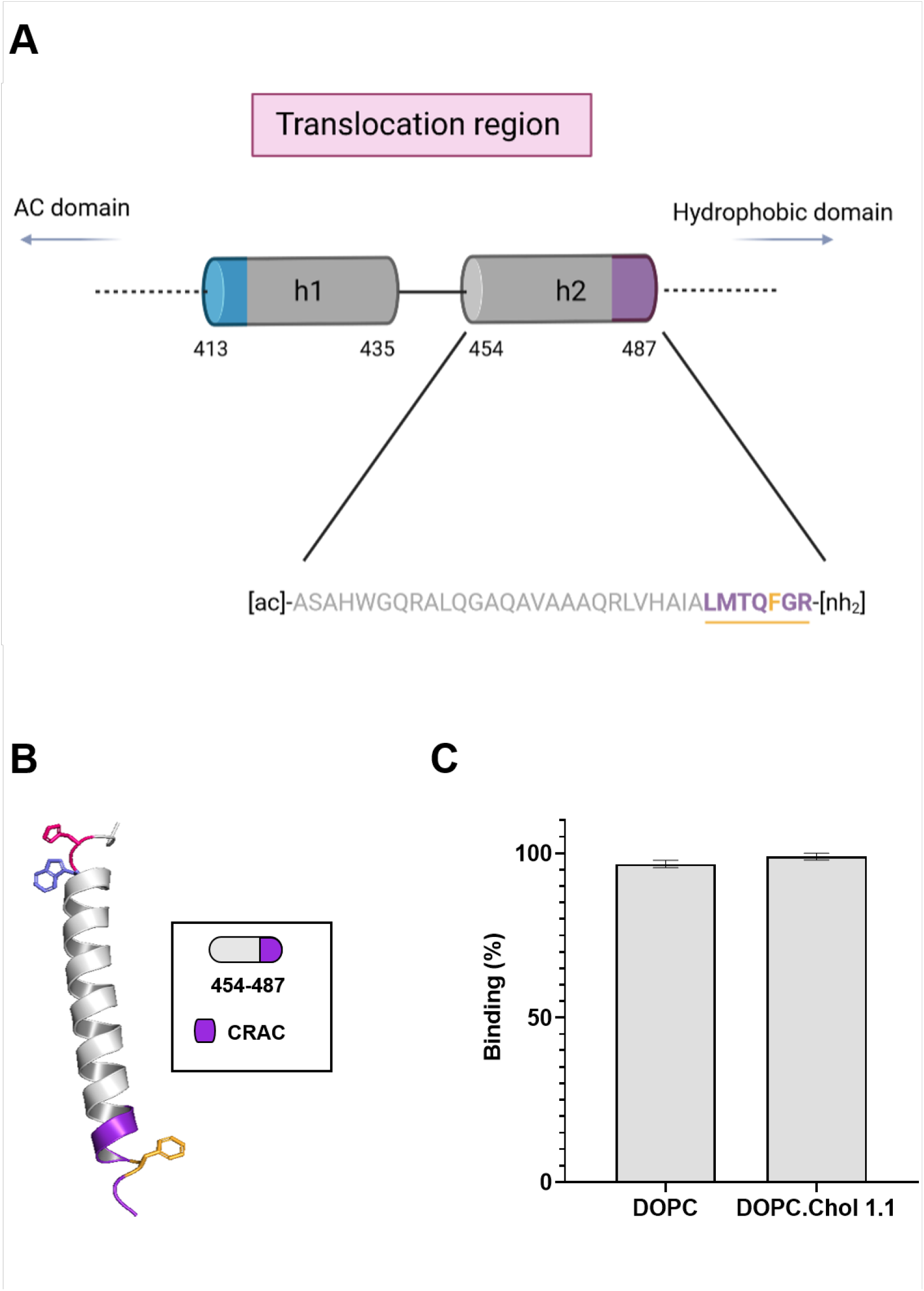
Binding of the P454-487 peptide to liposomes composed of pure DOPC or DOPC:Chol (1:1). (**a**) Schematic drawing of the ACT translocation region showing two constituting α-helices, h1 (residues 418 to 439) and h2 (residues 457 to 485) [39]. Sequence of the peptide P454-487 used in this study is also shown in the figure. At the C-terminal end of P454-487 sequence the CRAC motif (in purple color) with a central aromatic residue, Phe-485 (in orange color) is highlighted. Drawing done with BioRender. (**b**) Model of the most likely secondary structure adopted by P454-487 as predicted by AlphaFold Protein Structure Database. The C-terminal CRAC motif is indicated with purple color (**c**) Binding of the P454-487 peptide to liposomes composed of pure DOPC or DOPC:Chol (1:1 molar ratio). Two-way ANOVA and Tukeýs multiple comparison test were performed (ns p=0.8988).

### P454-485 conformation in DOPC lipid bilayers

To explore the molecular basis of the interaction of the P454-487 peptide with membranes we investigated whether addition of lipid membranes induced conformational changes in the peptide structure, and whether the lipid composition, particularly, the presence or not of cholesterol, could have any differential effect on the peptide conformation.

Secondary structure changes of P454-487 were followed by transmission FTIR (Fourier Transform Infra-Red) spectroscopy. Fig 2 displays the amide Í region of the IR spectrum of the peptide in solution, and the corresponding band decomposition of the spectrum, showing a prominent band centered at 1645 cm^-1^ (random coil), which accounted about the 62% of the total conformers in buffer. In contrast, upon peptide reconstitution in DOPC bilayers, predominant helical conformers amounted to ca. 90% (64% α-helix plus 26 % helix solvated). Besides, in comparison with the absorption band components measured in TFE (2,2,2-Trifluoroethanol) 20%, the contribution of the 1675 cm^−1^ band (turns) and random coils decreased in DOPC bilayers, whereas the amide-Í band became overall much narrower (Supplementary Figure 1). These spectral variations reflect a reduction in the conformational space accessible to the P454-487 peptide chain upon reconstitution in the DOPC lipid bilayers, consistent with the majority of the membrane-associated peptide adopting a canonical α-helical conformation.

**Figure 2.**
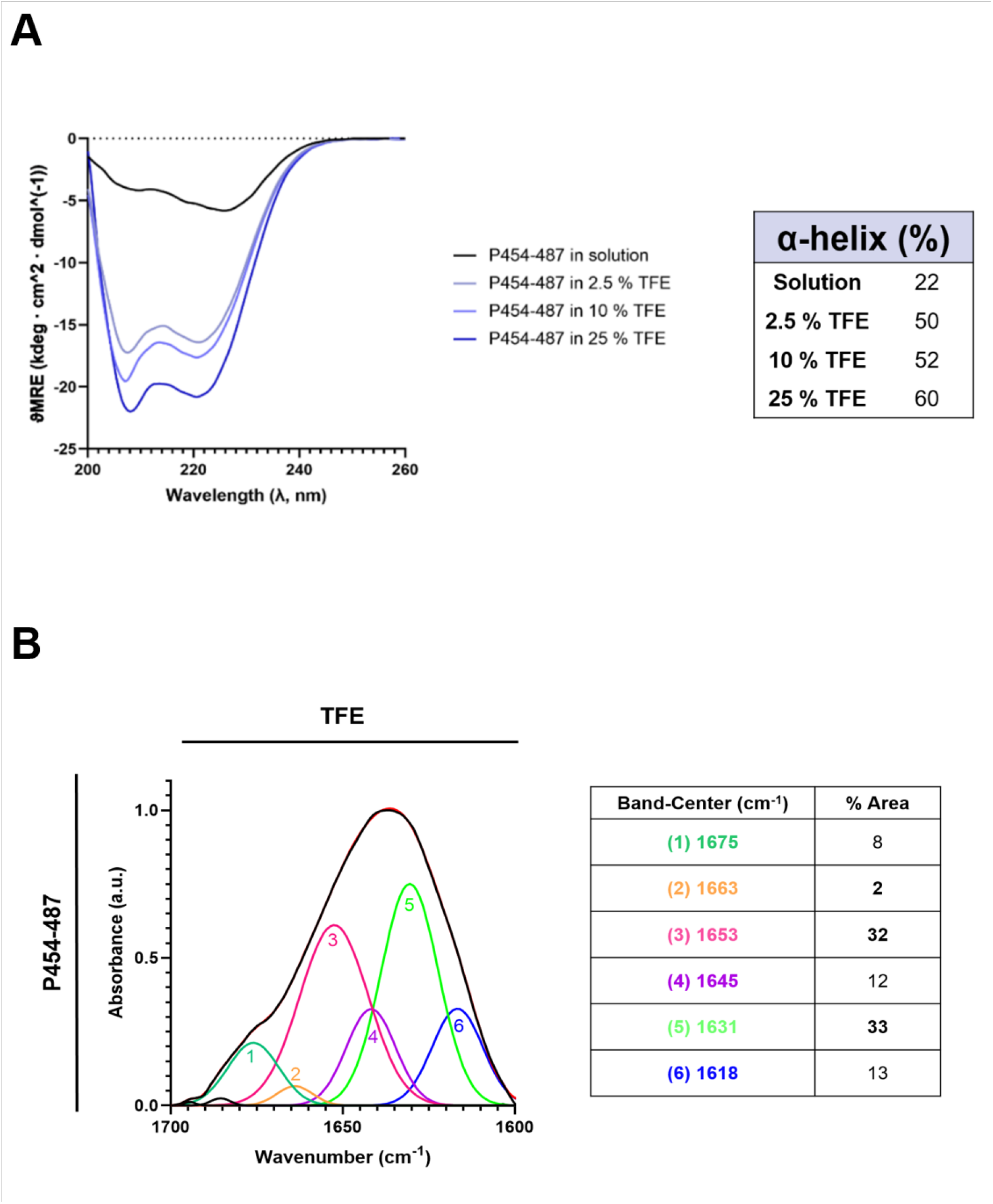
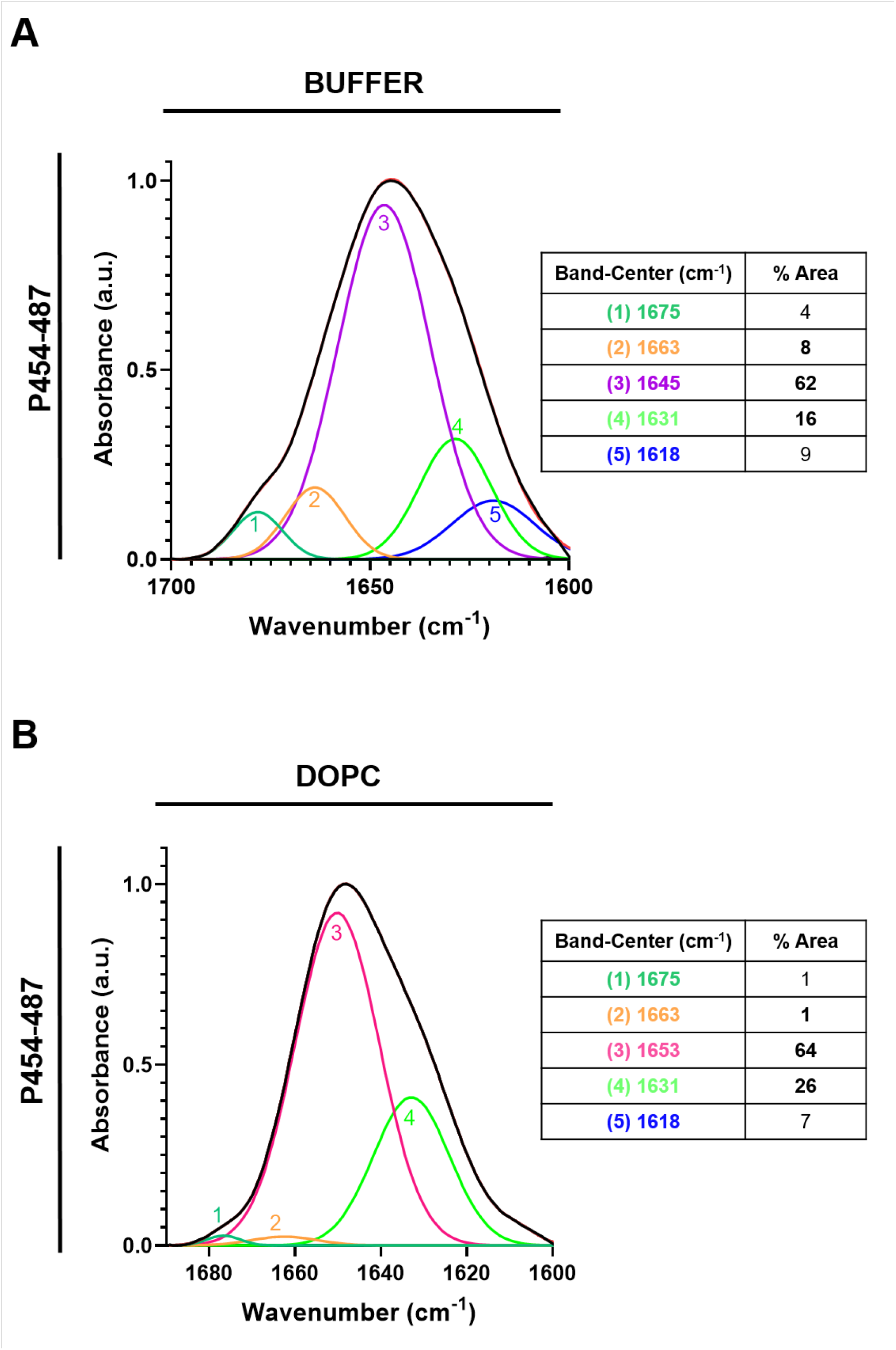
Normalized FT-IR spectra of the amide Í region for the P454-487 peptide in solution and reconstituted in DOPC lipid bilayers. (**a**) Normalized FT-IR spectrum of the amide Í region for the P454-487 peptide in solution. (**b**) Normalized FT-IR spectrum of the amide Í region for the P454-487 peptide reconstituted in DOPC bilayers. The absorption bands were decomposed into different components. The original spectra and the sum of the band components are superimposed and indistinguishable. The inset display the secondary structure assignation for the main components (bands labelled with numbers 1 to 5) and the area percentages (rounded off to the nearest integer).

### Conformational changes of P454-487 in cholesterol-containing membranes

To address whether the presence of cholesterol could have any differential effect on the P454-487 peptide conformation, we analyzed the conformation adopted by the peptide in membranes containing increasing cholesterol concentrations. Fig 3a displays the series of IR spectra as a function of cholesterol content in membranes and Table I summarizes the secondary structure assignation for the main components (bands labelled with numbers 1 to 6) and the corresponding area percentages.

**Figure 3.**
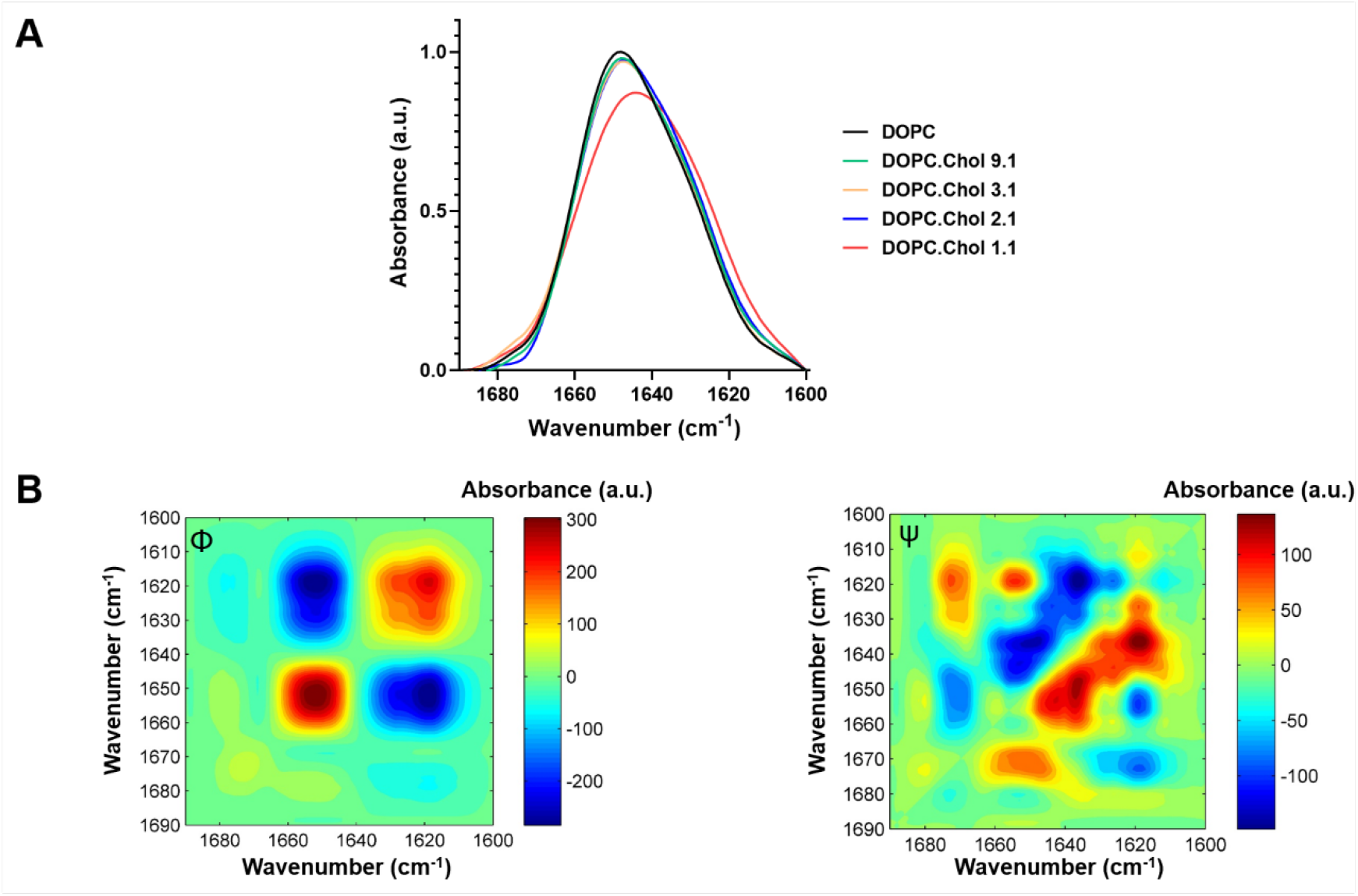
Conformations adopted by P454-487 as a function of the cholesterol content in membranes. (**a**) Normalized FT-IR spectra of the amide Í region of P454-487 reconstituted in DOPC bilayers with increasing cholesterol concentrations (peptide-to-lipid molar ratio, 1:50). **(b)** 2D-COS IR analysis of the IR spectra obtained for the P454-487 reconstituted in DOPC with increasing cholesterol concentrations. Synchronous (Φ, left) and Asynchronous (ψ, right) correlation map contours of the IR spectra obtained with increasing cholesterol concentrations are shown. Red peaks correspond to positive correlations and blue peaks to negative ones.

**Table I.** Secondary structure assignation for the main components of the decomposed spectra of P454-487 peptide in different environments and the area percentages for each one (rounded off the nearest integer).

| Band-Center | SOLUTION | 20 % TFE | DOPC | DOPC:Chol 9:1 | DOPC:Chol 3:1 | DOPC:Chol 2:1 | DOPC:Chol 1:1 |
| --- | --- | --- | --- | --- | --- | --- | --- |
| (1) 1675 | 5 | 9 | 1 | 1 | 1 | 1 | 0 |
| (2) 1663 | 17 | 1 | 1 | 0 | 0 | 0 | 2 |
| (3) 1653 | 0 | 33 | 66 | 71 | 72 | 51 | 49 |
| (4) 1645 | 57 | 14 | 0 | 0 | 0 | 0 | 12 |
| (5) 1631 | 10 | 30 | 25 | 24 | 25 | 46 | 22 |
| (6) 1618 | 11 | 13 | 6 | 2 | 0 | 1 | 15 |
| (7) 1606 | 0 | 0 | 1 | 1 | 2 | 1 | 0 |

Incorporation of low cholesterol concentrations (10-25%) into the DOPC lipid bilayers induced only modest changes in the P454-487 peptide conformation, causing a 6% increment of the α-helical conformation and the disappearance of the 1620 cm^-1^ band (extended conformation) (Fig 3a). The increase of the cholesterol content from 25% to 33 % induced instead a 16% decrease of the α-helix compared to pure DOPC. Moreover, the presence of the helix solvated was prominently increased, almost duplicating its contribution (from 26% to 46%). Interestingly, a further augmentation in the cholesterol concentration, from 33% to 50%, reverted the helix solvated component, which decreased to the half (from 46% to 22%), while the buried α-helical component remained practically unchanged. In addition, a new band corresponding to random coil appeared (12%) and the band corresponding to extended conformations increased to 15% (see Table I).

To get more insight into the P454-487 conformational changes induced by the membrane cholesterol concentration, we next performed the 2D-correlation analysis of the IR spectra in the corresponding amide Í band region [68-70]. We note that relevant effects detected on the 2D maps often reflect subtle changes in the relative contents of the amide I band components [68-70]. The analysis performed revealed the evolution of the spectral main components (Fig 3b).

In the synchronous (Φ) 2D map of the raw spectra of P454-487 (Fig 3b), the two auto-peaks indicated simultaneous changes in the bands composing the amide-Í spectrum. These two auto-peaks were found centered at 1653 and 1620 cm^−1^, whereas the two cross-relation negative peaks 1653/1620 cm^−1^ and 1620/1653 cm^−1^ reflected that both vibrations were affected in-phase by cholesterol, the first component (α-helix) diminishing in intensity as cholesterol in the membrane increases and the second (extended conformation) augmenting.

The corresponding asynchronous (Ψ) maps reflected the sequential order of events induced by the increase of cholesterol (Fig 3b). The asynchronous peaks were positive (red contours) if the change in the first frequency occurred accelerated with respect to that in the second one, and negative (blue contours) if delayed [68-70]. Several positive and negative correlation peaks are detected in the asynchronous (Ψ) map. As mentioned before, it is known that partial solvation of α-helical structures can give rise to low-frequency bands centered at ca. 1635–1630 cm^−1^ because of the cross-hydrogen bonds that can be formed with water [71, 72]. Thus, we have attributed the P454-487 absorption mode at 1635 cm^−1^ to a fraction of the helical structure not buried in the membrane, i.e., exposed to solvent and/or in contact with interfacial polar moieties. Accordingly, the positive correlation peak 1635/1655 cm^−1^ detected in the maps suggests the protrusion and exposition of part of the buried α-helix to a more polar solvated location, likely because the lipid bilayer becomes less and less fluid as the cholesterol concentration increases. The second major positive correlation peak 1620/1635 cm^−1^ suggests a conversion of the solvated helix fraction into an extended conformation.

In conclusion, the IR data support that the increment in the cholesterol content in the membrane up to 33% leads to the protrusion of part of the buried α-helix, which becomes exposed to solvent and/or in contact with interfacial polar moieties. The increment of the cholesterol concentration from 33% to 50% seems to promote then conversion of a fraction of the exposed helix into an extended conformation, and the unfolding of other fraction. Interestingly, even at the highest cholesterol concentration, the fraction of the buried α-helix (ca. 50%) remains being the predominant component of the P454-487 peptide conformations in the membranes.

### Membrane insertion angle of the P454-487 peptide

The previous results support the efficient reconstitution of the P454-487 peptide as α-helix in DOPC lipid bilayers, and the possibility that part of it transits to an extended structure in cholesterol-enriched membranes. Using ATR-IR spectroscopy (Atennuated Total Reflection IR), we next determined the tilt of the P454-487 peptide conformations relative to the membrane normal in the DOPC and DOPC:Chol 1:1 lipid bilayers. ATR-IR is a useful method for determining the orientation of α-helices of proteins or peptides in the lipid bilayer [73-76]. ATR-IR absorbance spectra were measured using perpendicular and parallel polarized light. From these spectra, the experimental average dichroic ratios were calculated, and order parameters *S* and tilt angles calculated (Supplementary Table 1) [73-76].

According to the tilt angle inferred from the dichroic ratios, the longitudinal axis of the P454-487 helix formed an angle of 52° with the DOPC lipid bilayer normal, and an angle of 49° with the DOPC:Chol (1:1) lipid bilayer normal, suggesting thus, that P454-487 may insert into the membrane in a similar tilted orientation in both lipid compositions.

### Membrane association of the P454-487-F485A mutant peptide to neutral lipid bilayers

In the past, other laboratory had studied the interaction with lipid bilayers of another synthetic peptide (so-called P454 by the authors) [77]. P454 is very similar to our P454-487 peptide, except by the absence of the last three C-terminal residues (485 to 487, FGR) that are present in P454-487 peptide. As reported by the authors P454 acquires α-helical secondary structure upon interaction with a lipid bilayer, but in contrast to the observed here for P454-487, P454 lies parallel to the membrane plane [77]. Furthermore, P454 associates to vesicles containing negatively charged phospholipids such as POPG or POPS inducing their permeabilization at peptide concentrations in the range of micro-molar. However, it does not partition to and permeabilize neutral liposomes [77]. It appears therefore that P454 peptide has rather different properties relative to the P454-487 peptide explored here. The interesting point is that both peptides differ only in the presence or absence of three C-terminal residues that are part of a functional CRAC motif identified in the full length ACT by our laboratory [38] (see Fig 1). P454-487 peptide contains thus an intact cholesterol recognition site (^481^LMTQFGR^487^), while P454 peptide has an incomplete CRAC motif (^481^LMTQ^484^). Consequently, we asked whether these three extra residues could confer different structural characteristics to P454-487 that might explain the different interaction mode of the peptide with lipid membranes, or whether the important point is to have or not an intact CRAC motif that can bind cholesterol.

To address this question, another peptide (P454-487-F485A) was synthesized in which the central Phe-485 residue was substituted by Ala. The aromatic residue Phe (or Tyr) is a key residue in the CRAC motifs [49, 50], and previously we had found that such substitution in the full length ACT inhibits both the lytic and translocation activities of the toxin [38].

We determined first the binding capacity of P454-487-F485A peptide to liposomes of DOPC or DOPC: Chol (1:1 molar ratio), finding a slightly lower peptide binding to the DOPC LUVs, and a significantly inferior association to the sterol-containing vesicles (Fig 4a). Given that the only difference between the two peptides was the Phe-485, we hypothesized that this residue may be relevant in the cholesterol recognition by the native P454-487 peptide.

**Figure 4.**
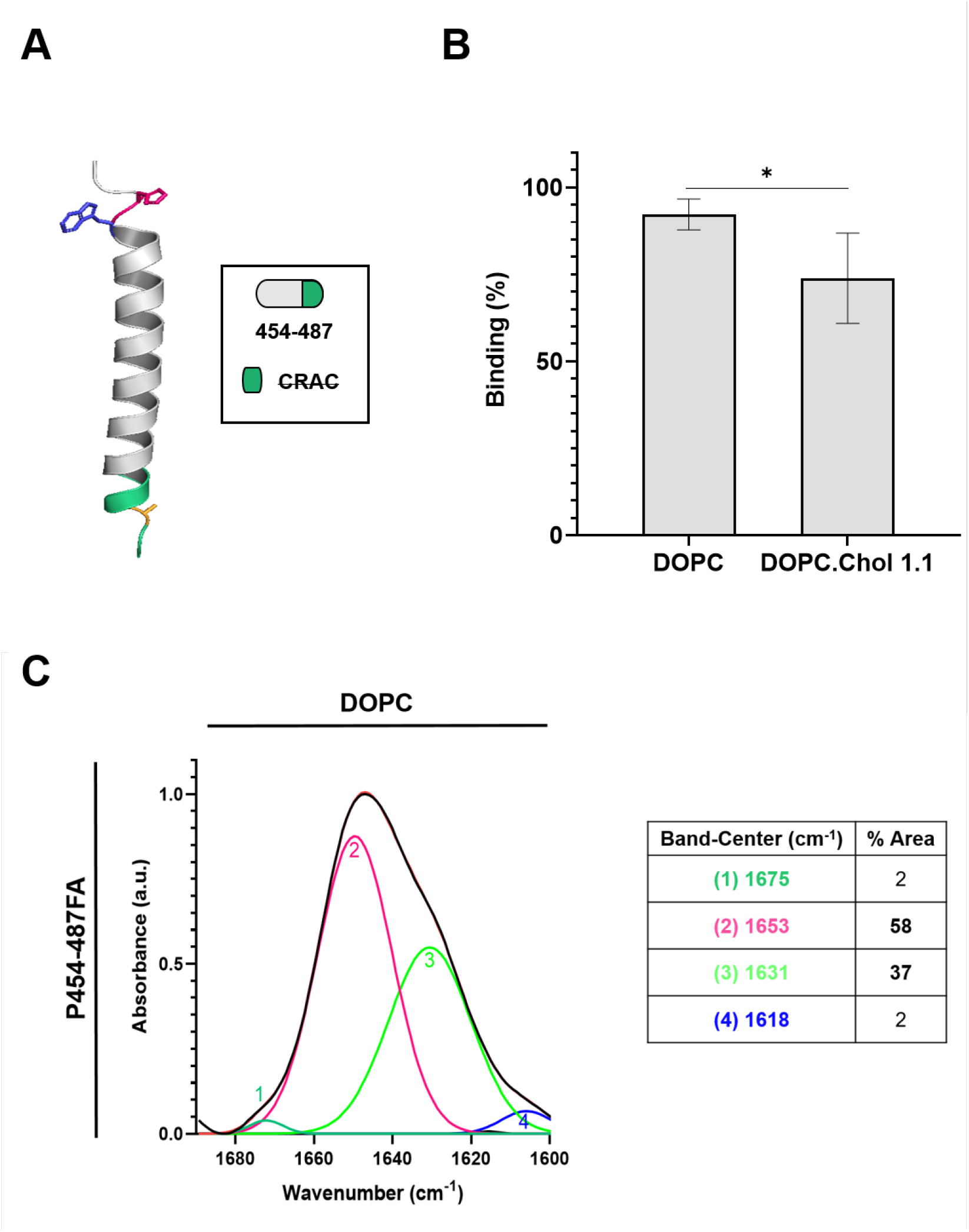
Binding of the P454-487F485A mutant peptide to liposomes composed of pure DOPC or DOPC:Chol (1:1 molar ratio). (**a**) Model of the most likely secondary structure adopted by the mutant peptide as determined by the AlphaFold Protein Structure Database. **(b)** Binding of the P454-487F485A peptide to liposomes composed of pure DOPC or DOPC:Chol (1:1 molar ratio). Two-way ANOVA and Tukeýs multiple comparison test were performed (* p=0.0325). **(c)** Normalized FT-IR spectrum of the amide Í region for the P454-487F485A peptide reconstituted in DOPC lipid bilayers. The absorption band was decomposed into different components. The original spectrum and the sum of the band components are superimposed and indistinguishable. The inset displays the secondary structure assignation for the main components (bands labelled with numbers 1 to 4) and the area percentages (rounded off to the nearest integer).

### Conformation adopted by the P454-487F485A mutant peptide in lipid bilayers

We explored first the conformation of P454-487F485A in DOPC bilayers. As shown in Fig 4b and Table II almost the 100% of the mutant peptide was helicoidal in these conditions. To note that the helix solvated fraction (ca 36%) resulted being greater than the determined for P454-487 reconstituted in liposomes of the same lipid composition (see Table II).

**Table II.** Secondary structure assignation for the main components of the decomposed spectra of P454-487F485A in different environments, and the area percentages for each one of the components (rounded off the nearest integer).

| Band-Center | SOLUTION | 20 % TFE | DOPC | DOPC:Chol 9:1 | DOPC:Chol 3:1 | DOPC:Chol 2:1 | DOPC:Chol 1:1 |
| --- | --- | --- | --- | --- | --- | --- | --- |
| (1) 1675 | 3 | 4 | 2 | 2 | 2 | 0 | 0 |
| (2) 1663 | 16 | 13 | 0 | 0 | 0 | 2 | 1 |
| (3) 1653 | 13 | 18 | 58 | 59 | 62 | 55 | 61 |
| (4) 1645 | 42 | 25 | 0 | 0 | 0 | 0 | 0 |
| (5) 1631 | 16 | 25 | 37 | 34 | 33 | 41 | 33 |
| (6) 1618 | 8 | 9 | 2 | 0 | 0 | 0 | 0 |
| (7) 1606 | 1 | 1 | 3 | 3 | 3 | 2 | 5 |

Next, it was assayed the effect of cholesterol in the secondary structure of P454-487F485A upon its reconstitution in liposomes with increasing cholesterol content (10% to 50%) (Fig 5a). In this case, addition of cholesterol, irrespective of its proportion, had no significant effect on the overall helical conformation of the mutant peptide. The proportion of the helix exposed to the solvent remained practically invariable for all the lipid compositions, representing about one third of the total (Table II). Interestingly, this solvated helix proportion was greater than the observed for the native peptide (≈25% for all the lipid mixtures), except for the DOPC:Chol 2:1 liposomes in which it was of about the 50% (Table II).

**Figure 5.**
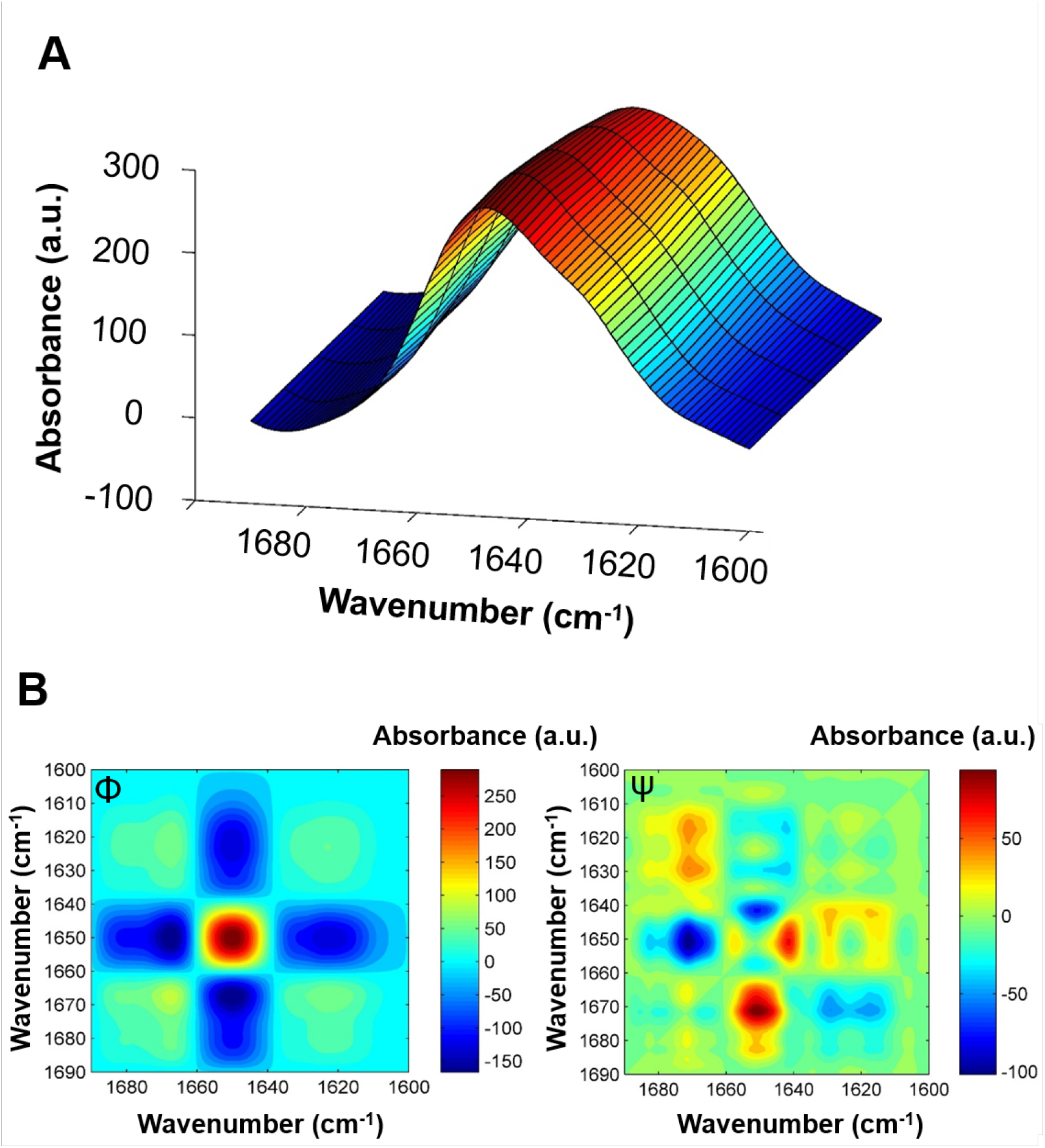
2D-COS IR analysis of the IR spectra obtained for the P454-487F485A reconstituted in DOPC with increasing cholesterol concentrations. (**a**) Normalized 3D FT-IR spectra of the amide Í region of P454-487F485A reconstituted in DOPC bilayers with increasing cholesterol concentrations (peptide-to-lipid molar ratio 1:50). (**b**) 2D-COS IR analysis of the IR spectra obtained for the P454-487F485A reconstituted in DOPC with increasing cholesterol concentrations. Synchronous (Φ, left) and asynchronous (Ψ, right) correlation map contours of the IR spectra obtained with increasing cholesterol concentrations. Red peaks correspond to positive correlations and blue peaks to negative ones.

The 2D-COS IR analysis of the IR spectra obtained for the P454-487F485A mutant peptide was in line with these findings (Fig 5b). The Synchronous map (Φ) of the P454-487F485A IR spectra showed auto-peaks in 1653 cm^-1^, band corresponding to the α-helix (Fig 5b). This might be related to subtle changes in the maximum intensity of this band, due to a small displacement of it.

In concordance, the Asynchronous map (ψ) also displays small changes (Fig 5b). Variation on the helix environment might alter the amide Í vibrational modes, even if the secondary structure does not change. Thereupon, FT-IR data and 2D-COS IR analysis support that the addition of cholesterol does not trigger structural interconversion of the P454-487F485A peptide, and so it can be affirmed that cholesterol does not have effect on the overall secondary structure adopted by this peptide in cholesterol-containing membranes.

Finally, using ATR-IR spectroscopy, we determined the tilt of the P454-487F485A peptide conformations relative to the membrane normal in the DOPC and DOPC:Chol lipid bilayers (Supplementary Table 2). In accordance with the tilt angle inferred from the dichroic ratios, the longitudinal axis of the P454-487F485A helix formed an angle of 43° with the DOPC lipid bilayer normal. Interestingly, this angle progressively increased from 43% to ca. 64% as the cholesterol content in the bilayer augmented, suggesting that the peptide was progressively expelled out from the bilayer interior as the bilayer was becoming more and more rigid and tightly packed.

Briefly, our data concur in that the mutant peptide is a long alpha helix, with about two thirds of it penetrating the hydrophobic part of the bilayer in a tilted orientation and the remaining third accessible to the solvent, and which responds to the presence of increasing concentrations of cholesterol simply by augmenting the tilt angle. This is consistent with an oblique peptide that inserts less and less deeply into the hydrophobic core as the lipid packing becomes more and more rigid in presence of cholesterol. We demonstrate thus that the membrane-interacting properties of the P454-487F485A and P454-487 peptides are clearly different.

### Inhibition of the ACT-induced RBC hemolysis and cAMP production by the P454-487 peptide

Having determined that P454-487 was able to bind efficiently to neutral lipid bilayers, we tested whether it was capable of hindering ACT binding to natural membranes, and concomitantly of reducing or blocking ACT activity on red blood cells and macrophages.

To this end, we performed two different types of experiments. One approach consisted in pre-incubating a given ACT concentration with different peptide concentrations in solution, subsequently adding the mixture of the corresponding cells (RBC or macrophages), and then following the hemolytic kinetics (in RBC) or the cAMP production (in macrophages). As depicted in Fig 6a, the ACT-induced hemolysis was reduced by P454-487 in a peptide concentration-dependent way, with a diminishment of the RBC lysis by more than 50 % for the highest peptide dose used in the pre-incubation. Similarly, when ACT and the peptide were added concomitantly to the J774A.1 cells, it was observed a prominent reduction in the ACT-induced cAMP production, which fell already to almost the half with the lowest peptide concentration used, furtherly diminishing with the higher peptide doses assayed (Fig 6c).

**Figure 6.**
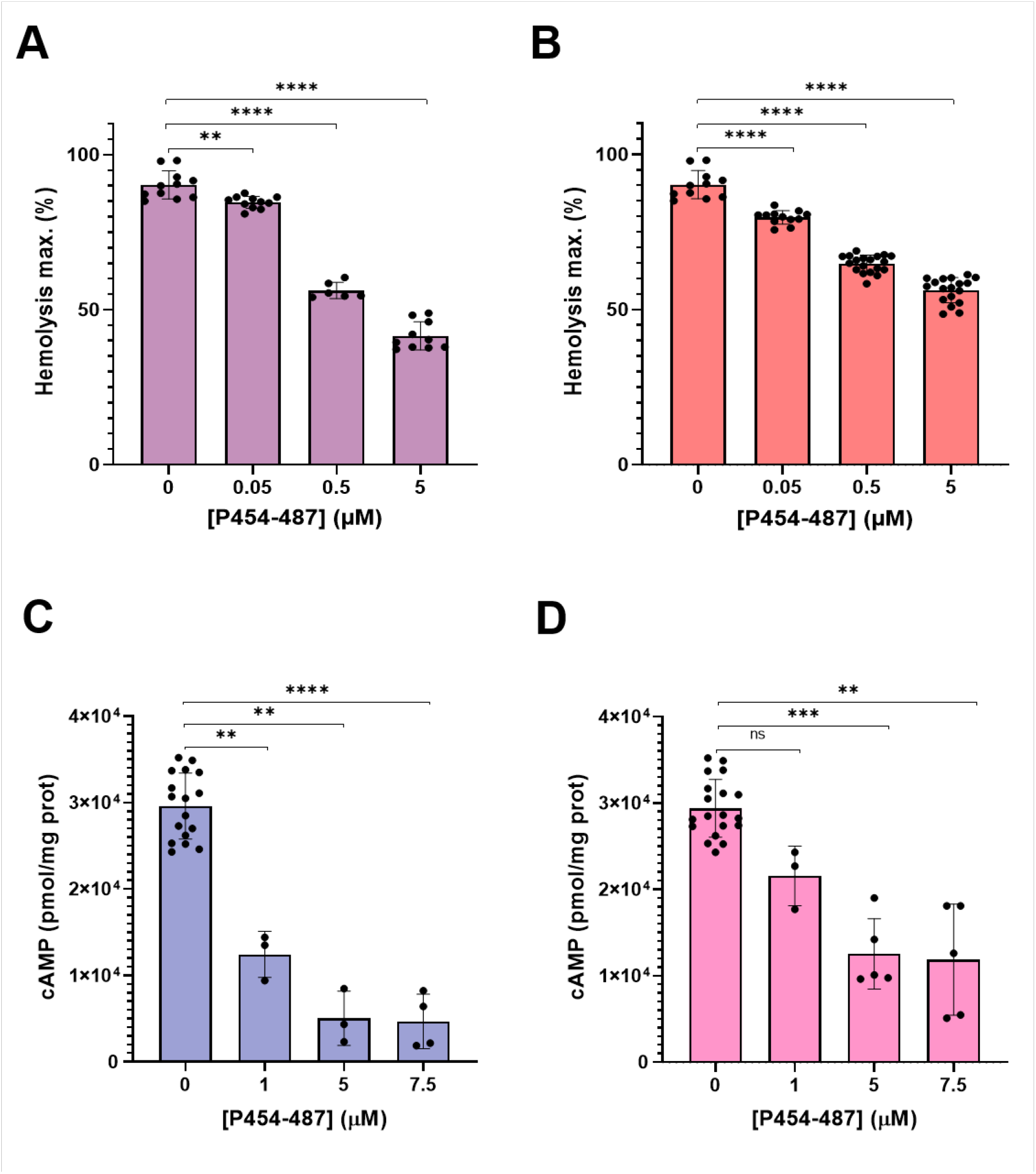
Effect of the pre-incubation of ACT with P454-487 on the toxin-induced RBC hemolysis and cAMP production in J774A.1 macrophages. (**a**) Maximum hemolysis caused by ACT (50 nM) on red blood cells (5 x 10^8^ cells/ml), after toxin pre-incubation with 0-5 µM P454-487 for 15 min. (**b**) Maximum hemolysis caused by ACT (50 nM) on red blood cells (5 x 10^8^ cells/ml), after pre-incubating red blood cells with 0-5 µM P454-487 for 15 min. In both cases, a further incubation of red blood cells was carried out for 180 min at 37°C. Brown-Forsythe and Welch’s ANOVA test was performed with Dunnet’s T3 multiple comparison test. ** p = 0.0032 and **** p < 0.0001 (**c**) cAMP production by ACT (1 nM) in J774A.1 macrophages (1 x 10^5^ cells/ml) after toxin pre-incubation for 10 min with different P454-487 concentrations (0–7.5 µM. (**d**) cAMP production by ACT (1 nM) in J774A.1 macrophages (1 x 10^5^ cells/ml) after pre-incubation of the macrophages with different P454-487 doses (0–7.5 µM) for 15 min. cAMP quantification was performed after incubation of the cells with ACT for 30 min at 37°C. Brown-Forsythe and Welch’s ANOVA test were performed with Dunnet’s T3 multìple comparison test.

In the second approach the cells (RBC or macrophages) were initially pre-incubated with different concentrations of the peptide, followed by the subsequent addition of ACT, and further incubation and recording of the hemolysis kinetics or cAMP determination (Figs 6b and 6d). In this case, it was also observed a peptide concentration-dependent decrease in the ACT-induced hemolysis, though the inhibitory effect exerted by the pre-bound peptide was more modest than in the first type of experiment (Fig 6a), likely because the peptide was not at saturating concentrations (Fig 6b). In the assay with the J774A.1 cells, a peptide-concentration-dependent inhibitory effect on the cAMP generation by ACT resulted very evident (Fig 6d). Together thus, these results corroborate that P454-487 is capable of hindering ACT association to the cell membrane of either RBC and macrophages, and consequently of hampering ACT-induced RBC lysis and cAMP accumulation in macrophages.

Finally, the ability of P454-487 itself to alter membrane permeability of red blood cells was examined, and compared with the lytic capacity of the full-length ACT toxin. It was important to determine if the CRAC peptide was itself toxic to cells, since this would prevent a future therapeutic use.

As compared to the full-length ACT, which provoked the lysis of erythrocytes at nM concentrations, no peptide-mediated hemolysis was observed in the treated erythrocytes, even upon incubation with high peptide concentrations up to 200 µM (Supplementary Figure 2).

## DISCUSSION

Binding and productive insertion into the lipid bilayer are the first essential stages in the action mechanism of any toxin acting at the level of cell membranes. Precluding this initial step may thus be a direct way to avoid their toxic effects on target cells.

It is known that plasma membrane cholesterol is essential for both the lytic and cytotoxic activity of *Bordetella* Adenylate Cyclase Toxin [38, 78]. Recently, we revealed that such cholesterol sensitivity is provided by four cholesterol-recognition motifs we identified in helical elements of the translocation region and pore-forming domain, since they are necessary for the two ACT activities [38]. We hypothesize that direct interaction with membrane cholesterol through those four cholesterol-binding motifs drives insertion and stabilizes the transmembrane topology of several helical elements in the TR and HR that ultimately build the ACT structure for AC delivery and pore-formation, thereby explaining the cholesterol-dependence of the ACT activities.

In the present study, we have explored whether a 34 residue-long synthetic peptide, containing at the C-terminus one of those four cholesterol-binding sites, is itself capable of binding to cell membranes, of inserting into the bilayer core, and, of avoiding ACT cytotoxicity on cells by outcompeting with the full-length toxin for the sterol binding. In addition, we have dissected the membrane-interacting properties of the peptide, and we discuss the relevance of our findings in the context of the full-length ACT.

We show that P454-487 associates effectively to membrane systems (large unilamellar liposomes) composed of neutral phospholipids (100% DOPC or DOPC: Chol 1:1), achieving almost 100% binding when assayed at a lipid: protein ratio of 500:1 (Fig 1). Further, we find that P454-487 peptide can efficiently outcompete with the full-length ACT for binding to the plasma membrane of target cells, which results in an effective, concentration-dependent inhibition of the ACT-induced hemolysis (Fig 6a,b) and of the cAMP accumulation in J774A.1 macrophages (Fig 7c,d). This reaffirms the high binding efficacy of P454-487 not only to nude pure lipid bilayers, but also to the more complex plasma membrane of living cells. Importantly, we show that P454-487 does not itself exhibit lytic properties in the range of concentrations in which inhibits ACT activity on cells (Supplementary Fig 1), all of which allows affirming that use of the P454-487 peptide may represent a valid non-toxic method for blocking binding of ACT to host cells and minimize ACT cytotoxicity.

Regarding the structure of the membrane-bound P454-487, we show that the membrane-associated peptide adopts a canonical α-helical conformation into pure DOPC bilayers, with about two thirds of the helix buried into the membrane core, while the remaining third is exposed to solvent and/or in contact with interfacial polar moieties (Fig 3 and Table I). This seems consistent with the great number of amino acid residues constituting the P454-487 peptide (34 amino acids), which exceed the ≈20 residues on average required by a transmembrane α-helix to span a phospholipid bilayer [79, 80]. Doing some simple calculations from these data, we could say that about 22 residues of the P454-487 helix would be spanning the DOPC bilayer, while other 12 residues would be outside the membrane. In addition, we find that the P454-487 helix penetrates the DOPC bilayer in a tilted orientation, forming an oblique angle of ≈52° relative to the membrane normal (Supplementary Table 1). Insertion with tilted orientation into phosphatidylcholine bilayers was reported in the past for single spanning transmembrane model peptides such as those of the WALP family [81] noting that tilt angles of transmembrane helixes are a compensation mechanism for hydrophobic mismatch. This is the difference between the thicknesses of hydrophobic regions of a transmembrane protein/peptide and of the lipid membrane it spans [82].

The finding of the modulating effect of cholesterol on the membrane-associated P454-487 conformation is also remarkable. We show that low cholesterol concentrations between 10 and 25% have little effect on the peptide conformation, causing only a small increment of ca. 6% in the α-helix component (Table I), which may be consistent with the increased thickness of the pure phospholipid bilayers in presence of cholesterol [83]. A greater sterol content (33 %) induces however a considerable increase in the solvent-exposed α-helix fraction, which almost duplicates the percentage in pure DOPC bilayers (Table I). This higher solvation of the helix may be most likely due to protrusion of part of the buried α-helix fraction, provoked by a more tightly packed bilayer in presence of the higher cholesterol concentration. This proves that P454-487 is sensitive to the lipid packing effect, which is fully coherent with part of the peptide penetrating into the more hydrophobic core formed by the acyl chains of DOPC. Interestingly, a further augmentation of the cholesterol concentration in the membrane, from 33% to 50%, provokes several conformational changes, but that apparently affect only to the protruded α-helix fraction of P454-487, which partly adopts β strand-like conformation (≈15%), while other part unfolds (≈12%) (Table I). Instead, the fraction of α-helix buried into the bilayer remains constant, representing ca. 50%. This suggests that even in tightly packed, cholesterol-rich, lipid bilayers P454-487 retains within the bilayer at least ≈17 of its 34 amino acid residues, which might be enough for the buried α-helix to span the lipid bilayer. This is striking if one considers that this sterol percentage (≈ 50%) is very similar to the amount present in the plasma membrane of cells such as erythrocytes and macrophages, where ACT is fully active, and might be of great relevance in the context of the full-length ACT. It might guarantee that the translocation region remains anchored within the lipid bilayer in an ample range of cholesterol concentrations. Finally, we show that in the different DOPC:Chol bilayers the buried fraction of the P454-487 helix also penetrates obliquely into the bilayer with an angle of 49°-51° relative to the membrane normal, very similar to the determined for the pure DOPC bilayers (Supplementary Table 1). Nevertheless we cannot exclude that in the context of the full-length ACT the tilt angle varies as has been described for helices in polytopic membrane proteins which have their tilt and rotation fully constrained by their tertiary structure [84].

Other peptides that were reported to penetrate cholesterol-rich lipid membranes in a tilted orientation and with similar oblique angles (45° - 50°) are the fusion peptides of the HIV-1 or Ebola viruses and peptides from various pathogenic proteins (α-synuclein or β-amyloid peptide) [85, 86]. These tilted peptides act as topological binding sites for cholesterol [85], but there is not particular consensus sequence characterizing a cholesterol-binding site on a tilted peptide. Rather, it is envisioned that the tilted orientation allows an excellent adaptation of the peptide to the inverted cone shape of cholesterol [86]. Thus, the capacity to interact with cholesterol appears to be intrinsic to tilted peptides.

A pertinent question is then, what role, if any, plays the CRAC motif present at C-terminus of P454-487? The answer is provided by P454-487F485A, the point mutant peptide. We find that cholesterol in the lipid bilayer, irrespective of its proportion, does not affect the overall secondary structure adopted by the P454-487F485A in membranes (Table II). This mutant peptide inserts, either in DOPC as in the different DOPC:Chol mixtures assayed, as a tilted α- helix, with about 60% of it buried into the lipid bilayer, and 40% remaining solvent accessible (Figs 5-6 and Table II). The only effect cholesterol seems to have on P454-487F485A is the progressive augmentation of the tilting angle of the buried helix, from 42° in the most fluid bilayer (DOPC), to ≈64° in a highly ordered packed membrane (DOPC:Chol 1:1, passing through 53° in a less ordered bilayer (DOPC:Chol 2:1) (Supplementary Table 2). Given that the only difference between P454-487 and P454-487F485A peptides is the defective CRAC-485 motif (Phe485Ala substitution) in the last one, we can conclude that presence of this intact cholesterol-recognition motif is relevant for the peptide-membrane association as may determine the membrane-anchoring properties of the P454-487 peptide in cholesterol-enriched membranes. We hypothesize that the conformational change experimented by the solvent accessible fraction of the P454-487 helix in cholesterol-rich membranes would allow to strengthen the peptide-cholesterol interaction, and/or to stablish new additional interactions likely with the polar heads of phospholipids, with the final outcome of retaining large part of the P454-487 α-helix buried within the bilayer (irreversible insertion). Going a step further, in the context of the full-length ACT molecule, the irreversible insertion of a given helix in the translocation region could furtherly contribute to determine the membrane topology of the neighboring helices in the TR and HD. The present data are thus fully consistent with previous results from our laboratory showing that the point mutation Phe485Ala in the CRAC-485 motif of the full-length ACT affects the lytic and translocation activities of the toxin [38]. They further support our hypothesis that a direct ACT interaction with membrane cholesterol through the four cholesterol-binding motifs in the TR and HD would stabilize the transmembrane topology of several helical elements that ultimately build the ACT structure for AC delivery and pore-formation [38]. Also consistently with our data others previously noted for another cholesterol-dependent RTX toxin (LtxA from *A. actinomycetemcomitans* leukotoxin), that the secondary structure of a short peptide containing a functional CRAC site changes from α- to β-structure as the cholesterol content of the bilayer increases up to 40% [87].

It is worth mentioning here, the recent study by Voegele and cols (2021) [88], in which the authors postulate that after ACT binding to target cells, the P454 segment (residues 454-484) destabilizes the plasma membrane, translocates across the lipid bilayer and binds calmodulin. They further propose that such calmodulin-P454 interaction in the cytosol may assist the transport of the 40 KDa N-terminal catalytic domain to the cytosol [88]. Here the authors are partly extrapolating previous conclusions of that the ACT segment extending between residues 454-484 exhibits membrane-active properties related to antimicrobial peptides to build their translocation model. The authors investigated the membrane-interacting properties of the synthetic peptide P454 (residues 454-484) [77], very similar to our P454-487 peptide, except by the absence in their peptide of the last three C-terminal residues (FGR, residues 485 to 487 in ACT) that are present in P454-487. They reported that P454 peptide associates only to vesicles containing negatively charged phospholipids, inducing their permeabilization at rather high peptide concentrations (µM range) and that acquires α-helical secondary structure upon interaction with a lipid bilayer, but in contrast to the observed here for the P454-487 peptide, P454 lies parallel to the membrane plane, [77]. It seems thus, that P454 peptide has rather different properties relative to the P454-487 peptide explored here, and remarkably, those differences seem to depend only on three residues, FGR, the absence of which conditions having an intact CRAC-485 motif. These data, thus, clearly reinforce the conclusion that the Phe-485 residue and intact C-terminal CRAC-485 motif critically determine the membrane-interacting properties of the 454-487 segment of the ACT TR region, while undermining the credibility of the P454-based translocation model, proposed in the aforementioned publication [88].

## METHODS

### Chemicals

Cholesterol (Chol) was purchased from Avanti Polar Lipids (Alabaster, AL) and 1,2-Dioleoyl-*sn*-glycero-3-phosphocholine (DOPC) from Sigma-Aldrich (St. Louis, MO).

### Peptide synthesis

The synthetic peptides P454-487 and P454-487FA were purchased from Genosphere Biotech (France) and their purity (99 %) and composition were controlled by reverse-phase HPLC and MALDI-mass spectrometry. The peptides are capped on the N-terminus with an acetyl group and on the C-terminus with an amide group. The P454-487 peptide has 34 amino acids and corresponds to residues 454-487 of ACT primary structure. Its amino acid sequence is the following: [ac]-ASAHWGQRALQGAQAVAAAQRLVHAIA**LMTQF^485^GR**-[NH_2_] (CRAC motif in bold). It has a molecular mass of 3628.14 g/mol, an estimated pI = 14, and an estimated charge of z = +3.2e at pH 7.00 (https://pepcalc.com/). Phe-485 was mutated to Ala in P454-487FA peptide (in red) and thus, its sequence is: [ac]-ASAHWGQRALQGAQAVAAAQRLVHAIA**LMTQA^485^GR**-[NH_2_]. Peptide stocks were stored in ethanol or MQ H_2_O, depending on the assay (See Supplementary Table SI).

### Expression and purification of ACT

ACT was expressed in Escherichia coli XL-1 blue cells (Stratagene) transformed with pT7CACT1 plasmid, kindly provided by Dr. Peter Sebo (Institute of Microbiology of the ASCR, v.v.i., Prague, Czech Republic) and purified as described by Karst et al. [56].

### Preparation of Lipid Vesicles

Large Unilamellar Vesicles (LUVs) were prepared at 10 mM concentrations from DOPC and DOPC:Chol 1:1 in liposome buffer (150 mM NaCl and 20 mM Tris, pH 8.00). Lipids dissolved in chloroform:methanol 2:1 (vol:vol) were added to a glass vial in the required amounts. Solvent was evaporated under a stream of nitrogen, and the residual was removed under vacuum, creating a thin lipid film on the glass surface. Multilamellar liposomes (MLV) were obtained by hydrating the lipid film with liposome buffer. LUVs were formed by extruding the MLV liposome solutions through a 100-nm polycarbonate membrane. The suspensions of LUVs were monodisperse with mean hydrodynamic diameters of 100 nm, as measured by dynamic light scattering on a NanoZS instrument (Malvern).

### Peptide reconstitution into lipid bilayers

To prepare peptide-containing vesicles, adequate amounts of lipids and peptide were mixed in organic solvent prior to the production of the liposomes as described [57]. Briefly, phospholipid and cholesterol were dissolved in chloroform:methanol 2:1 (vol:vol) and mixed with each peptide (dissolved in 100 % ethanol) at a peptide-to-lipid molar ratio of 1:50. The mixture was dried under a N_2_ stream followed by 2 h vacuum pumping to remove traces of organic solvents. Subsequently, the dried lipid films were hydrated with infrared buffer (150 mM NaCl, 20 mM Hepes, pH 7.4) at 55°C. Next, the multilamellar vesicles were bath sonicated (30min, RT) and subjected to 15 freeze and thaw cycles to obtain unilamellar vesicles. Same procedure was used for control samples that contained no lipid: peptide in aquous solution and peptide in 20 % TFE.

### Peptide binding assay

To measure binding of P454-487 and P454-487FA to membranes, a centrifugation assay was performed [58]. 20 µM peptide was incubated with DOPC or DOPC:Chol 1:1 liposomes (peptide:lipid ratio of 1:500), for 30 min at RT. Samples were added to a centrifugal filter (Amicon_ 30k MWCO; EMD Millipore, Billerica, MA) and centrifuged for 1 h at 6000 x g. Unbound peptide concentrations were calculated by comparing the intrinsic fluorescence of the eluate at 340 nm to a set of standards of the same peptide with known concentrations. The fluorescence measurements were recorded on a Fluorolog-3 spectrofluorometer (Horiba, Japan) using an excitation wavelength of 281 nm. The bound peptide concentrations were then calculated from the total and free concentrations of peptide.

### Circular dichroism (CD)

Circular dichroism (CD) measurements were carried out on a thermally-controlled Jasco J-810 circular dichroism spectropolarimeter calibrated routinely with (1 S)- (+)−10-camphorsulfonic acid, ammonium salt. 100 µM peptides were dissolved in an aqueous buffer (150 mM NaCl, 20 mM Tris, pH, 8.0) wiith increasing concentrations of TFE. Spectra were measured in a 1 mm path-length quartz cell equilibrated at 25 °C. Data were taken with a 1 nm band-width, 100 nm/min speed, and the results of 40 scans per sample were averaged. Contribution of the solvent was subtracted during data processing. The following (Eq. 1) was used to calculate alpha-helix content:

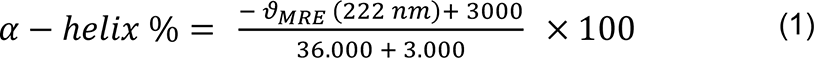

Where, *ϑ_MRE_* is calculated using (Eq. 2):

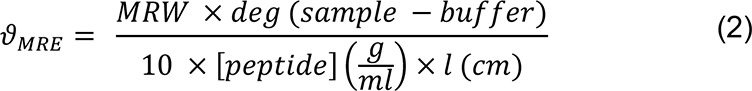

MRW is the Mean Residue Weight as defined by (Eq. 3):

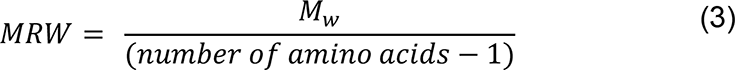

### Transmission infrared spectroscopy (FT-IR)

Secondary structure changes were followed by FT-IR spectroscopy. Peptide-containing samples (in absence and presence of lipid bilayers) were lyophilized and subsequently prepared at 2 mg (peptide)/mL in ²H₂O buffer. A 25 μl sample aliquot was deposited on a CaF_2_ window (BioCell, BioTools Inc., Wauconda, IL). Reference windows without peptide were prepared similarly. Infrared spectra were recorded in a Thermo Nicolet Nexus 5700 (Thermo Fisher Scientific; Waltham, MA) spectrometer equipped with a mercury-cadmium-telluride detector. Typically, 100 scans were collected for each background and sample, and the spectra were obtained with a nominal resolution of 2 cm^−1^. Data treatment and band decomposition of the original amide Í have been described elsewhere [59]. In brief, the number and position of bands were obtained from the deconvolved (bandwidth = 18 and k = 2) spectra. The baseline was removed before starting the fitting procedure and initial heights set at 90 % of those in the original spectrum for the bands in the wings and for the most intense component, and at 70 % of the original intensity for the rest of bands. An iterative process followed, in two stages: (i) The band position of the component bands was fixed, allowing widths and heights to approach final values; (ii) band positions were left to change. For band shape a Gaussian function was used. The restrictions in the iterative procedure were needed because initial width and height parameters can be far away from the final result due to the overlapping of bands, so that spurious results can be produced. In this way, information from band position, percentage of amide Í band area and bandwidth were obtained for every component. Using this procedure the result was repetitive. Mathematical accuracy was assured by constructing an artificial curve with the parameters obtained and subjecting it to the same procedure again. The number of bands was fixed on the basis of the narrowing procedures.

### Attenuated total reflection IR spectroscopy (ATR-IR)

ATR-IR spectra were measured in a Bruker Tensor 27 spectrometer (Bruker MA, USA) equipped with a mercury-cadmium-telluride detector using a BioATRCell II micro-ATR unit. 20 μl of the lipid mixtures containing peptide were dried on the surface of the ATR Ge crystal by flowing dried air into the infrared spectrometer chamber during 5 h. For spectra acquisition, the polarized mirror (Pike Technologies, WI, USA) was adjusted to 0° and 90°, to generate incident light oriented parallel and perpendicular to the lipid normal, respectively. 100 IR spectra at 2 cm^−1^ resolution were collected under each condition and averaged. The dichroic ratio of the amide Í bond absorption was computed for parallel (0°) versus perpendicular (90°) polarized incident light relative to the membrane normal and was employed to calculate the peptide orientation as discussed previously [60-62].

### Hemolysis assays

Hemolysis assays were performed on 96-well plates. First, we tested the lytic capacity of the peptides alone. For that purpose, an erythrocyte suspension (≈5 x 10^8^ cells/ml) was incubated with different peptide or ACT concentrations (0-200 µM or 0-400 nM respectively) for 180 min at 37°C. Samples were centrifuged then and supernatant absorbance was measured at 412 nm.

For competition assays, 50 nM ACT was pre-incubated with 0, 50, 500 or 5000 nM peptide in assay buffer (150 mM NaCl, 20 mM Tris pH 8.0, 2 mM CaCl_2_) at 4 °C for 15 min. In parallel, 50 nM ACT was preincubated with 0, 50, 500 and 5000 nM peptide for 15 min at 4 °C. In both cases, after the incubation time, these samples were added onto an equal volume of erythrocytes at a density of 5 x 10^8^ cells/ml. 180 min kinetics were acquired at 700 nm, at 37°C and constant stirring. The blank (0 % hemolysis) corresponded to erythrocytes incubated in buffer without toxin and 100 % hemolysis was obtained by adding Triton X-100 (0.1 %) to the erythrocyte suspension.

### Cell culture

J774A.1 macrophages (ATTC, number TIB-67) were grown at 37°C in DMEM containing 10 % (v/v) FBS, and 4 mM L-glutamine in a humidified atmosphere with 5 % CO_2_.

### Measurement of cAMP

cAMP assays were performed on 96-well plates. cAMP produced in cells was measured upon incubation of 1 nM ACT with J774A.1 cells (8 x 10^4^ cells/ ml) for 30 min at 37°C. In one set of experiments, cells were first pre-incubated with 0.0-7.5 µM peptides for 10 min. On other set, 1 nM ACT was pre-incubated with 0.0-7.5 µM peptides for 10 min. Then the mixture was added onto the cells for further incubation.

In both cases, 96-well plates were centrifuged at 4 °C, 600 x g, 10 min. Supernatant was removed and a solution of 0.1 M HCl and 0.3 % Triton X-100 was added for 10 min. Samples were frozen and thawed and centrifuged again. This time, supernatants were stored for the direct cAMP ELISA kit (Enzo Lifesciences).

## Acknowledgements

This study was funded by the Spanish Ministerio de Economía y Competitividad (grant number BFU2017–82758-P). J.A. was recipient of a fellowship from the University of Basque Country (UPV/EHU). RA holds a contract funded by the Fundación Biofisika Bizkaia.

## Author contributions

J.A., R.A. and H.O. conceived and designed the experiments; J.A. and R.A. performed experiments and analysed the data; I.d.l.A. and J.A. prepared samples and determined structures by FT-IR. H.O. wrote the paper with contributions from all the authors. All authors have read and agreed to the published version of the manuscript.

## Additional information

### Supplementary information

Correspondence and requests for materials should be addressed to H. Ostolaza.

## Supplementary Material for the manuscript

**Title**: Relevance of an intact cholesterol-recognition motif for the membrane-interacting properties of a peptide derived from the translocation region of the Adenylate Cyclase Toxin

**Authors**: Jone Amuategi, Rocío Alonso, Igor de la Arada & Helena Ostolaza

**Suplementary Figure 1.**
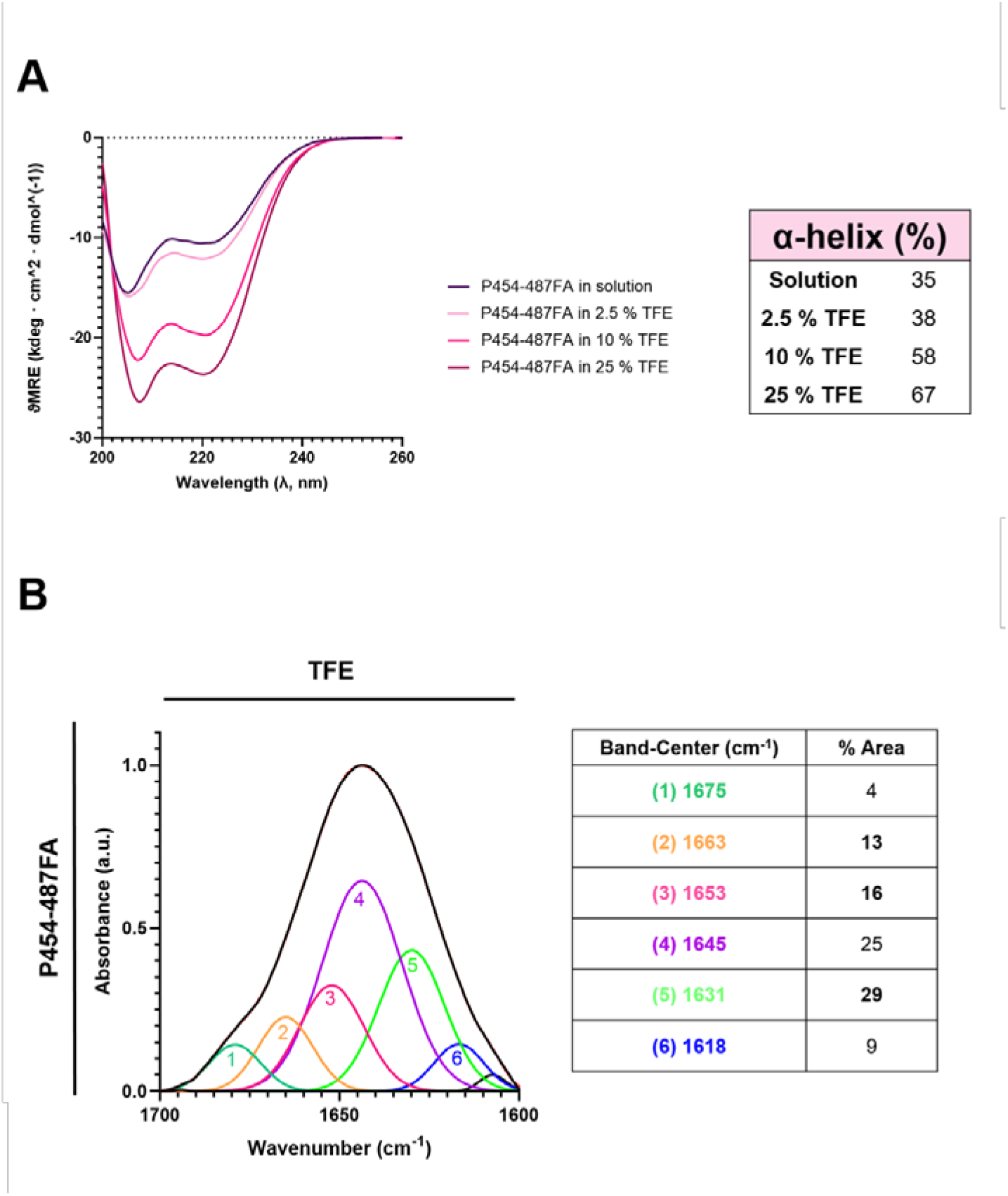
Secondary structure of the P454-487F485A peptide in TFE nonpolar medium. (**A**) CD spectra obtained for P454-487F485A at 25 °C in the presence of increasing concentrations of TFE, a medium that mimics the low-polarity of lipid bilayers. Content of α-helix (%) is represented for each condition. (**B**) FT-IR spectrum of the amide Í region for the P454-487F485A peptide in buffer containing 20 % TFE. The absorption band was decomposed into different components. The original spectrum and the sum of the band components are superimposed and indistinguishable. The inset displays the secondary structure assignation for the main components (bands labelled with numbers 1 to 6) and the area percentages.

**Suplementary Figure 2.**
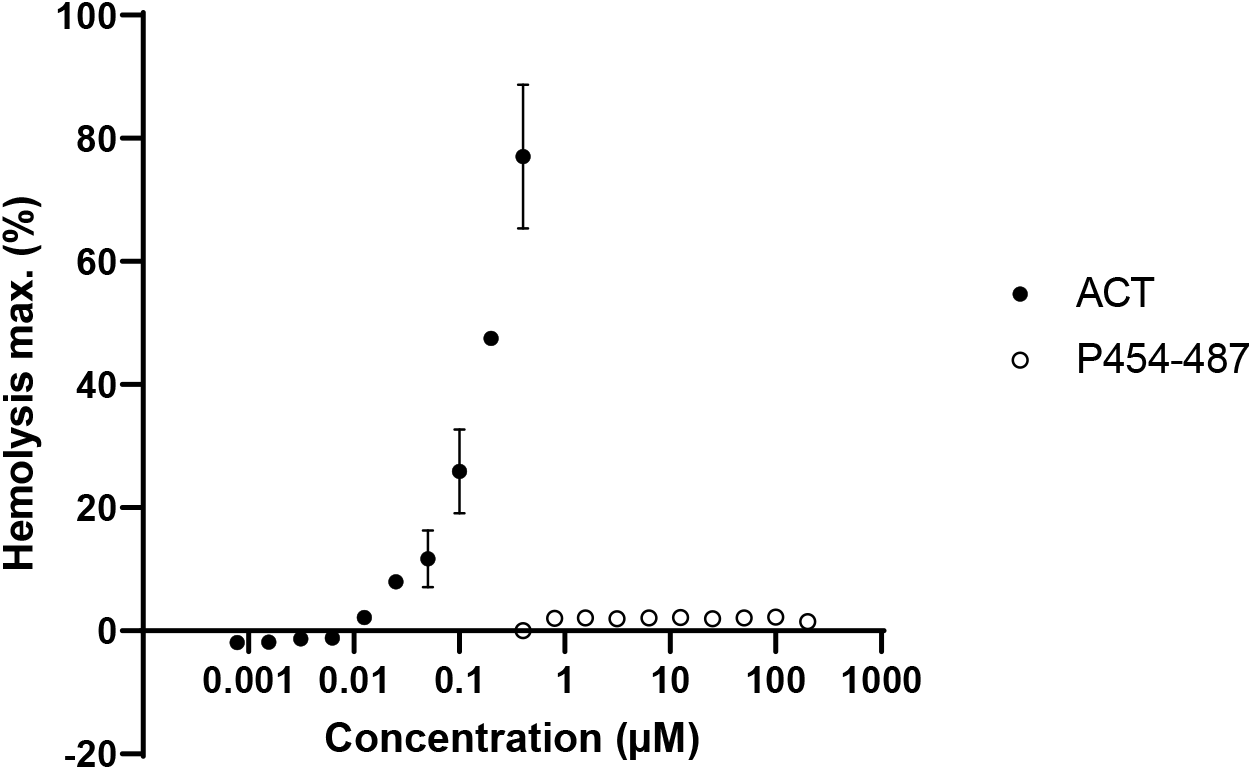
Lytic capacity of the P454-487 peptide on red blood cells and comparison with the full-length ACT toxin. Maximum hemolysis caused by ACT (0-400 nM) or P454-487 (0-200 μM) on red blood cells (5 x 10^8^ cells/ml), after incubating the samples at 37 °C for 180 min.

**Supplementary Table I.**
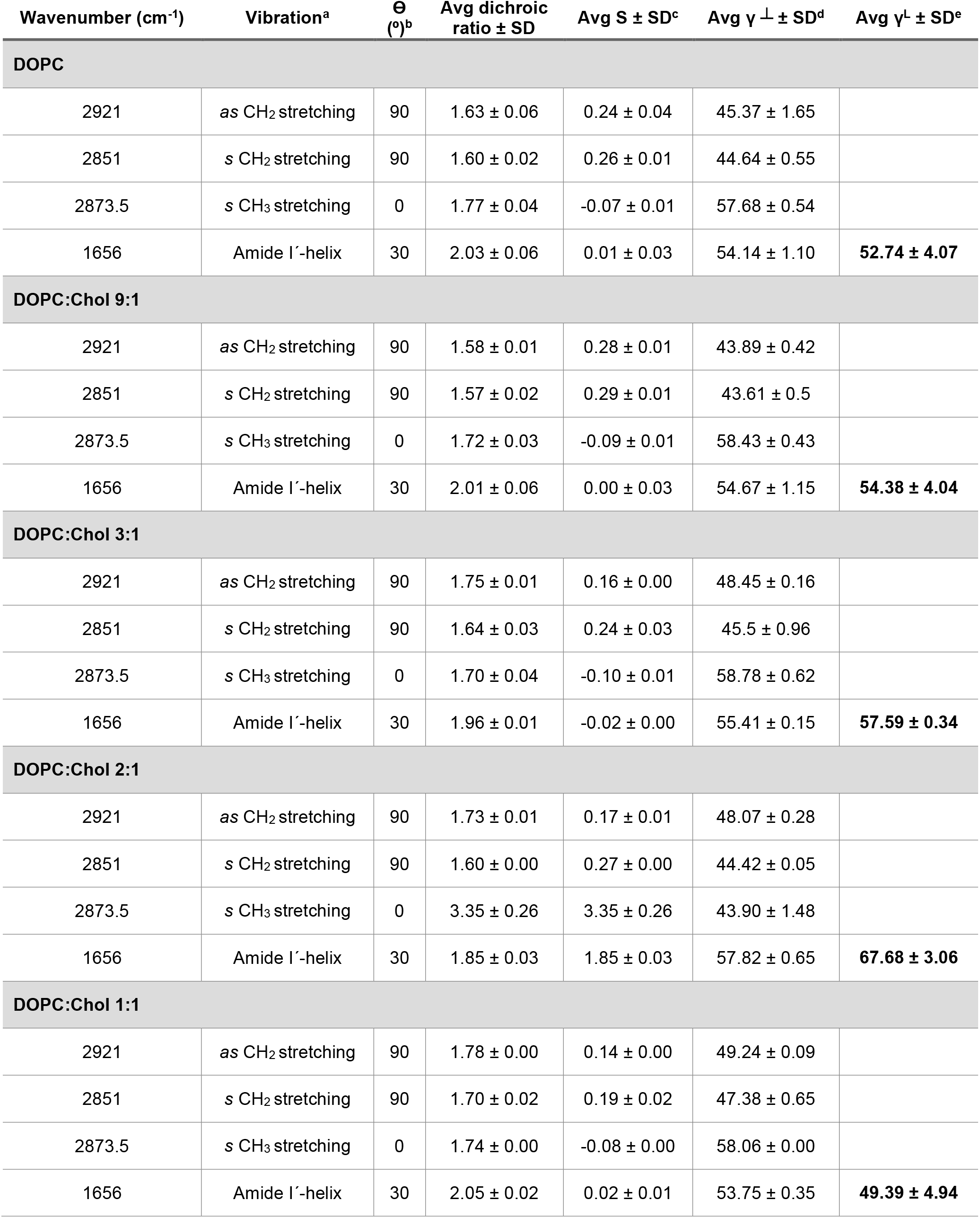
ATR-IR data of the P454-487 peptide. ^a^Vibrations are presented as symmetric (*s*) or asymmetric (*as*). ^b^, direction of the dipole moment associated with the vibration with respect to the direction of the main molecular axis (aliphatic chain or peptide-secondary structure). ^c^S, form factor. ^d^ γ ┴, angle between the direction of the molecular axis and the perpendicular to the crystal plane (similar to the membrane plane). ^e^γ^L^, angle between the direction of the peptide-secondary structure axis and the calculated aliphatic chain axis.

**Suplementary Table II.**
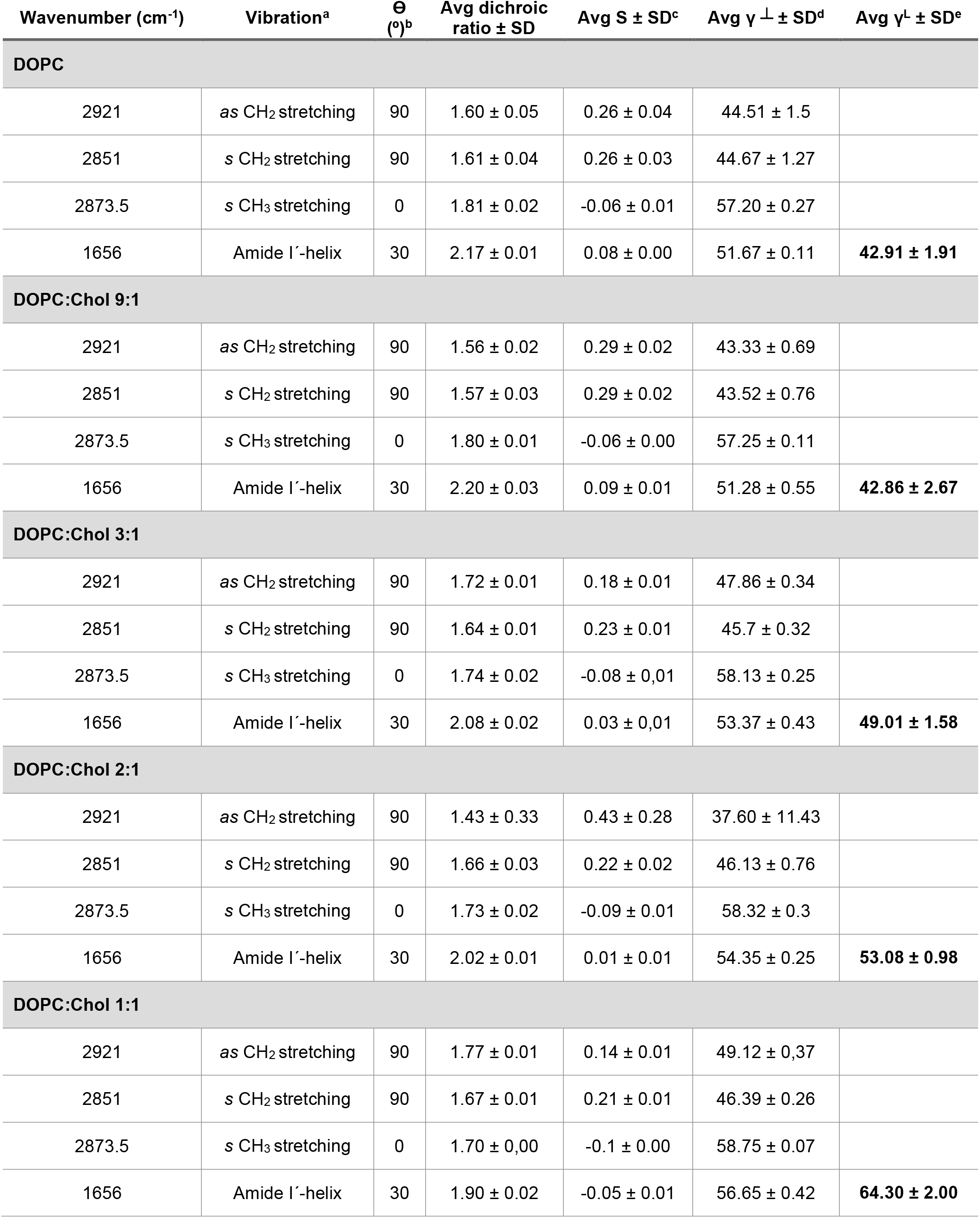
ATR-IR data of the P454-487FA peptide. ^a^Vibrations are presented as symmetric (*s*) or asymmetric (*as*). ^b^, direction of the dipole moment associated with the vibration with respect to the direction of the main molecular axis (aliphatic chain or peptide-secondary structure). ^c^S, form factor. ^d^ γ ┴, angle between the direction of the molecular axis and the perpendicular to the crystal plane (similar to the membrane plane). ^e^γ^L^, angle between the direction of the peptide-secondary structure axis and the calculated aliphatic chain axis.

## REFERENCES

[1] Carbonetti NH. Immunomodulation in the pathogenesis of *Bordetella pertussis* infection and disease. Current Opinion in Pharmacology 7, 272–278 (2007)

[2] Hewlett, E.L. Whooping Cough and Other *Bordetella* Infections. Goldman’s Cecil Medicine: 24th Edition 2, 1900–1903 (2012)

[3] Mattoo, S, Cherry, JD. Molecular pathogenesis, epidemiology, and clinical manifestations of respiratory infections due to *Bordetella pertussis* and other *Bordetella* subspecies. Clin. Microbiol. Rev. 18, 326–382 (2005)

[4] Melvin JA, Scheller, EV, Miller, JF, Cotter, PA. Bordetella pertussis pathogenesis: current and future challenges. Nat Rev Microbiol. 12, 274–288 (2014)

[5] Sealey KL, Belcher T, Preston. *Bordetella pertussis* epidemiology and evolution in the light of pertussis resurgence. Infect Genet Evol. 40, 136-143 (2016)

[6] Saeidpour A, Bansal S, Rohani P. Dissecting recurrent waves of pertussis across the boroughs of London. PLoS Comput Biol. 18, e1009898 (2022)

[7] Linhartová I., Bumba L., Mašín J., Basler M., Osička R., Kamanová J., Procházková K., Adkins I., Hejnová-Holubová J., Sadílková L., Morová J., Sebo P. RTX proteins. A highly diverse family secreted by a common mechanism. FEMS Microbiol. Rev. 34, 1076–1112 (2010)

[8] Welch R. A. Pore-forming cytolysins of Gram-negative bacteria. Mol. Microbiol. 5, 521–528 (1991)

[9] Coote J. G. Structural and functional relationships among the RTX toxin determinants of Gram-negative bacteria. FEMS Microbiol. Rev. 8, 137–161 (1992)

[10] Welch, R A. RTX toxin structure and function: a story of numerous anomalies and few analogies in toxin biology. Curr. Top. Microbiol. Immunol. 257, 85–111 (2001)

[11] Gray, M.C., Lee, S.-J., Gray, L.S., Szabo, G., Hewlett, E.L. Translocation-specific conformation of adenylate cyclase toxin from Bordetella pertussis inhibits toxin-mediated hemolysis. Journal of Bacteriology 183, 5904–5910 (2001)

[12] Karst J. C., Barker R., Devi U., Swann M. J., Davi M., Roser S. J., Ladant D., Chenal A. Identification of a region that assists membrane insertion and translocation of the catalytic domain of *Bordetella pertussis* CyaA toxin. J. Biol. Chem. 287, 9200–9212. (2012)

[13] Osičková, A., Osička, R., Maier, E., Benz, R., Šebo, P. An amphipathic α-helix including glutamates 509 and 516 is crucial for membrane translocation of adenylate cyclase toxin and modulates formation and cation selectivity of its membrane channels. J. Biol. Chem. 274, 37644–37650 (1999)

[14] Benz, R., Maier, E., Ladant, D., Ullmann, A., Sebo, P. Adenylate cyclase toxin (CyaA) of *Bordetella pertussis*. Evidence for the formation of small ion-permeable channels and comparison with HlyA of Escherichia coli. J. Biol. Chem. 269, 27231–27239 (1994)

[15] Hackett, M., Guo, L., Shabanowitz, J., Hunt, D.F., Hewlett, E.L. Internal lysine palmitoylation in adenylate cyclase toxin from *Bordetella pertussis*. Science, 266, 433–435 (1994)

[16] Westrop G. D., Hormozi E. K., Da Costa N. A., Parton R., Coote J. G. *Bordetella pertussis* adenylate cyclase toxin. ProCyaA and CyaC proteins synthesised separately in *Escherichia coli* produce active toxin *in vitro*. Gene 180, 91–99 (1996)

[17] Bauche C., Chenal A., Knapp O., Bodenreider C., Benz R., Chaffotte A., Ladant D. Structural and functional characterization of an essential RTX subdomain of *Bordetella pertussis* adenylate cyclase toxin. J. Biol. Chem. 281, 16914–16926 (2006)

[18] Rose T., Sebo P., Bellalou J., Ladant D. Interaction of calcium with *Bordetella pertussis* adenylate cyclase toxin. Characterization of multiple calcium-binding sites and calcium-induced conformational changes. J. Biol. Chem. 270, 26370–26376 (1995)

[19] Guermonprez P., Khelef N., Blouin E., Rieu P., Ricciardi-Castagnoli P., Guiso N., Ladant D., Leclerc C. The adenylate cyclase toxin of *Bordetella pertussis* binds to target cells via the α(M)β(2) integrin (CD11b/CD18). J. Exp. Med. 193, 1035–1044 (2001)

[20] El-Azami-El-Idrissi, M., Bauche, C., Loucka, J., …Ladant, D., Leclerc, C. Interaction of *Bordetella pertussis* adenylate cyclase with CD11b/CD18. Role of toxin acylation and identification of the main integrin interaction domain. J. Biol. Chem. 278, 38514–38521 (2003)

[21] Glaser, P., Sakamoto, H., Bellalou, J., Ullmann, A., Danchin, A. Secretion of cyclolysin, the calmodulin-sensitive adenylate cyclase-haemolysin bifunctional protein of *Bordetella pertussis*. EMBO J 7, 3997-4004 (1988)

[22] Thomas, S, Holland, IB, Schmitt, L. The type 1 secretion pathway. The hemolysin system and beyond. Biochim Biophys Acta 1843, 1629-1641 (2014)

[23] Gao, Z., Young, R.A., Trucco, M.M., …Matschinsky, F.M., Wolf, B.A. Protein kinase A translocation and insulin secretion in pancreatic β-cells: Studies with adenylate cyclase toxin from *Bordetella pertussis*. Biochemical Journal 368, 397–404 (2002)

[24] Eby, J.C., Ciesla, W.P., Hamman, W., …Hewlett, E.L., Lencer, W.I. Selective translocation of the *Bordetella pertussis* adenylate cyclase toxin across the basolateral membranes of polarized epithelial cells. J. Biol. Chem. 285, 10662–10670 (2010)

[25] Hasan S, Kulkarni NN, Asbjarnarson A, Linhartova I, Osicka R, Sebo P, Gudmundsson GH. *Bordetella pertussis* Adenylate Cyclase Toxin Disrupts Functional Integrity of Bronchial Epithelial Layers. Infect Immun. 86, e00445-17 (2018)

[26] Martín, C., Requero, M.-A., Masin, J., …Sebo, P., Ostolaza, H. Membrane restructuring by *Bordetella pertussis* adenylate cyclase toxin, a member of the RTX toxin family. Journal of Bacteriology 186, 3760–3765 (2004)

[27] Wolff J., Cook G.H., Goldhammer A.R., Berkowitz S.A. Calmodulin activates prokaryotic adenylate cyclase. Proc. Natl. Acad. Sci. USA. 77, 3841–3844 (1980)

[28] Confer D.L., Eaton J.W. Phagocyte impotence caused by an invasive bacterial adenylate cyclase. Science 217, 948–950 (1982)

[29] Ehrmann, IE, Gray, MC, Gordon, VM, Gray, LS, Hewlett, EL Hemolytic activity of adenylate cyclase toxin from *Bordetella pertussis*. FEBS Lett. 278, 79-83 (1991)

[30] Gray, M., Szabo, G., Otero, A.S., Gray, L., Hewlett, E. Distinct mechanisms for K^+^ efflux, intoxication, and hemolysis by *Bordetella pertussis* AC toxin. J. Biol. Chem. 273, 18260–18267 (1998)

[31] González-Bullón, D., Uribe, K.B., Largo, E., …Martín, C., Ostolaza, H. Membrane permeabilization by *Bordetella* adenylate cyclase toxin involves pores of tunable size. Biomolecules 9, 183 (2019)

[32] Koji Tanaka, Jose M.M. Caaveiro, Koldo Morante, Juan Manuel González-Mañas, Kouhei Tsumoto. Structural basis for self-assembly of a cytolytic pore lined by protein and lipid. Nat Commun. 6, 6337 (2015)

[33] Bakrac B., et al. Molecular determinants of sphingomyelin specificity of a eukaryotic pore-forming toxin. J. Biol. Chem. 283, 18665–18677 (2008)

[34] Johnstone BA, Joseph R, Christie MP, Morton CJ, McGuiness C, Walsh JC, Böcking T, Tweten RK, Parker MW. Cholesterol-dependent cytolysins: The outstanding questions. IUBMB Life 74, 1169-1179 (2022)

[35] Brown, A.C., Balashova, N.V., Epand, R.M., …Boesze-Battaglia, K., Lally, E.T. *Aggregatibacter actinomycetemcomitans* leukotoxin utilizes a cholesterol recognition/amino acid consensus site for membrane association. J. Biol. Chem, 288, 23607–23621 (2013)

[36] Vazquez, R.F., Maté, S.M., Bakás, L.S., …Malchiodi, E.L., Herlax, V.S. Novel evidence for the specific interaction between cholesterol and α-haemolysin of *Escherichia coli*. Biochemical Journal 458, 481–489 (2014)

[37] Osickova, A., Balashova, N., Masin, J., …Lally, E.T., Osicka, R. Cytotoxic activity of *Kingella kingae* RtxA toxin depends on post-translational acylation of lysine residues and cholesterol binding. Emerging Microbes and Infections 7, 178 (2018)

[38] Amuategi, J., Alonso, R., Ostolaza, H. The role of four cholesterol-recognition motifs localized between amino acid residues 400-550 in regulating translocation and lytic activity of Adenylate Cyclase Toxin. Int J Mol Sci. 23, 8703 (2022)

[39] Sukova A., Bumba L., Srb P., Ververka V., Stanek O., Holulova J., Chmelik J., Sebo P., Masin J. Negative charge of the AC-to-Hly linking segment modulates calcium-dependent membrane activities of Bordetella adenylate cyclase toxin. Biochimica et Biophys. Acta Biomembr. 1862, 18310–18316 (2020)

[40] Masin J., Osickova A., Sukova A., Fiser R., Halada P., Bumba L., Linhartova I., Osicka R., Sebo P. Negatively charged residues of the segment linking the enzyme and cytolysin moieties restrict the membrane-permeabilizing capacity of adenylate cyclase toxin. Sci. Rep. 6, 29137 (2016)

[41] Prangkio P., Juntapremjit S., Koehler M., Hinterdorfer P., Angsuthanasombt C. Contributions of the hydrophobic Helix 2 of the *Bordetella pertussis* CyaA-hemolysin to membrane permeabilization. Protein Pept. Lett. 25, 236–243 (2018)

[42] Roderova J., Osickova A., Sukova A., Mikusova G., Fiser R., Sebo P., Osicka R., Masin J. Residues 529 to 549 participate in membrane penetration and pore-forming activity of the *Bordetella* adenylate cyclase toxin. Sci. Rep. 9, 5758 (2019)

[43] Osickova A., Osicka R., Maier E., Benz R., Sebo P. An amphipathic alpha-helix including glutamates 509 and 516 is crucial for membrane translocation of adenylate cyclase toxin and modulates formation and cation selectivity of its membrane channels. J. Biol. Chem. 274, 37644–37650 (1999)

[44] Basler M., Knapp O., Masin J., Fiser R., Maier E., Benz R., Sebo P., Osicka R. Segments crucial for membrane translocation and pore-forming activity of *Bordetella* adenylate cyclase toxin. J. Biol. Chem. 282, 12419–12429 (2007)

[45] Powthongchin B., Angsuthanasombat C. Effects on haemolytic activity of single proline substitutions in the *Bordetella pertussis* CyaA pore-forming fragment. Arch. Microbiol. 191, 1–9 (2009)

[46] Juntapremjit S., Thamwiriyasati N., Kurehong C., Prangkio P., Shank L., Powthongchin B., Angsuthanasombat C. Functional importance of the Gly cluster in transmembrane helix 2 of the *Bordetella pertussis* CyaA-hemolysin: Implications for toxin oligomerization and pore formation. Toxicon 106, 14–19 (2015)

[47] Li H, Papadopoulos V. Peripheral-type benzodiazepine receptor function in cholesterol transport. Identification of a putative cholesterol recognition/interaction amino acid sequence and consensus pattern. Endocrinology 139, 4991–4997 (1998)

[48] Fantini J & Barrantes. How cholesterol interacts with membrane proteins: an exploration of cholesterol-binding sites including CRAC, CARC and tilted domains. Front Physiol. 4, 31-41 (2013)

[49] Jacques Fantini, Coralie Di Scala, Luke S. Evans, Philip T. F. Williamson, Francisco J. Barrantes A mirror code for protein-cholesterol interactions in the two leaflets of biological membranes. Sci Rep. 6, 21907 (2016)

[50] Jafurulla M, Tiwari S, Chattopadhyay A. Identification of cholesterol recognition amino acid consensus (CRAC) motif in G-protein coupled receptors. Biochem Biophys Res Commun. 404, 569–573 (2011)

[51] Baier, C.J., Fantini, J., Barrantes, F.J. Disclosure of cholesterol recognition motifs in transmembrane domains of the human nicotinic acetylcholine receptor. Scientific Reports 1, 69 (2011)

[52] Boesze-Battaglia K, Brown A, Walker L, Besack D, Zekavat A, Wrenn S, Krummenacher C, Shenker BJ. Cytolethal distending toxin-induced cell cycle arrest of lymphocytes is dependent upon recognition and binding to cholesterol. J Biol Chem. 284, 10650–10658 (2009)

[53] Greenwood AI, Pan J, Mills TT, Nagle JF, Epand RM, Tristram-Nagle S. CRAC motif peptide of the HIV-1 gp41 protein thins SOPC membranes and interacts with cholesterol. Biochim Biophys Acta, Biomembr. 1778, 1120–1130 (2008)

[54] Vishwanathan SA, Thomas A, Brasseur R, Epand RF, Hunter E, Epand RM. Hydrophobic substitutions in the first residue of the CRAC segment of the gp41 protein of HIV. Biochemistry 47,124–130 (2008)

[55] Thaa B, Levental I, Herrmann A, Veit M. Intrinsic membrane association of the cytoplasmic tail of influenza virus M2 protein and lateral membrane sorting regulated by cholesterol binding and palmitoylation. Biochem J. 437, 389–397 (2011)

[56] Johanna C. Karst, V. Yvette Ntsogo Enguéné, Sara E. Cannella, Orso Subrini, Audrey Hessel, Sylvain Debard, Daniel Ladant, Alexandre Chenal. Calcium, Acylation, and Molecular Confinement Favor Folding of *Bordetella pertussis* Adenylate Cyclase CyaA Toxin into a Monomeric and Cytotoxic Form. J Biol Chem. 289, 30702–30716 (2014)

[57] Apellaniz, B. & Nieva, J. L. Fusion-competent state induced by a C-terminal HIV-1 fusion peptide in cholesterol-rich membranes. Biochim. Biophys. Acta 1848, 1014–1022 (2015)

[58] Angela C. Brown, Evan Koufos, Nataliya Balashova, Kathleen Boesze-Battaglia, Edward T. Lally. Inhibition of LtxA Toxicity by Blocking Cholesterol Binding With Peptides. Mol Oral Microbiol. 31, 94–105 (2016)

[59] Arrondo, J. L. & Goni, F. M. Structure and dynamics of membrane proteins as studied by infrared spectroscopy. Prog. Biophys. Mol. Biol. 72, 367–405 (1999)

[60] Menikh, A., Saleh, M. T., Gariepy, J. & Boggs, J. M. Orientation in lipid bilayers of a synthetic peptide representing the C-terminus of the A1 domain of shiga toxin. A polarized ATR-FTIR study. Biochemistry 36, 15865–15872 (1997)

[61] Marsh, D. Dichroic ratios in polarized Fourier transform infrared for nonaxial symmetry of beta-sheet structures. Biophys. J. 72, 2710–2718 (1997)

[62] Goormaghtigh, E., Raussens, V. & Ruysschaert, J. M. Attenuated total reflection infrared spectroscopy of proteins and lipids in biological membranes. Biochim. Biophys. Acta 1422, 105–185 (1999)

[63] Storrs, R.W., Truckses, D. and Wemmer, D.E. Helix propagation in trifluoroethanol Biopolymers 32,1695-1702 (1992)

[64] Blanco FJ, Jiménez MA, Pineda A, Rico M, Santoro J, Nieto JL. NMR solution structure of the isolated N-terminal fragment of protein-G B1 domain. Evidence of trifluoroethanol induced native-like beta-hairpin formation. Biochemistry 33, 6004–6014 (1994)

[65] Dyson HJ, Merutka G, Waltho JP, Lerner RA, Wright PE. Folding of peptide fragments comprising the complete sequence of proteins. Models for initiation of protein folding. I. Myohemerythrin. J Mol Biol. 226, 795–817 (1992)

[66] Whitmore L, Wallace BA. Protein secondary structure analyses from circular dichroism spectroscopy: methods and reference databases. Biopolymer 89, 392–400 (2008)

[67] Whitmore L, Wallace BA. DICHROWEB, an online server for protein secondary structure analyses from circular dichroism spectroscopic data. Nucleic Acids Res. 32, W668–W673 (2004)

[68] Fabian H, Mantsch HH, Schultz CP. Two-dimensional IR correlation spectroscopy: sequential events in the unfolding process of the lambda cro-V55C repressor protein. Proc. Natl. Acad. Sci. USA. 96, 13153–13158 (1999)

[69] Noda I. Two-dimensional correlation analysis useful for spectroscopy, chromatography, and other analytical measurements. Anal. Sci. 23,139–146(2007)

[70] Iloro I, Chehin R, Goni FM, Pajares MA, Arrondo JL. Methionine adenosyltransferase alpha-helix structure unfolds at lower temperatures than beta-sheet: a 2D-IR study. Biophys. J. 86, 3951–3958 (2004)

[71] Walsh ST, Cheng RP, Wright WW, Alonso DO, Daggett V, Vanderkooi JM, DeGrado WF. The hydration of amides in helices; a comprehensive picture from molecular dynamics, IR, and NMR. Protein Sci. 12, 520-531 (2003)

[72] Vu DM, Myers JK, Oas TG, Dyer RB. Probing the folding and unfolding dynamics of secondary and tertiary structures in a three-helix bundle protein. Biochemistry 43, 3582-3589 (2004)

[73] Menestrina G. Use of Fourier-transformed infrared spectroscopy for secondary structure determination of staphylococcal pore-forming toxins. Methods Mol. Biol. 145, 115–132 (2000)

[74] Vandenbussche, G., A. Clerckx, M. Clerckx, J. Curstedt, J. Johansson, H. Jörnvall, and J. M. Ruysschaert. Secondary structure and orientation of the surfactant protein SP-B in a lipid environment. A FTIR spectroscopy study. Biochemistry 31, 9169–9176 (1992)

[75] Menikh A, Saleh MT, Gariepy J, Boggs JM. Orientation in lipid bilayers of a synthetic peptide representing the C-terminus of the A1 domain of shiga toxin. A polarized ATR-FTIR study. Biochemistry 36, 15865–15872 (1997)

[76] Shai Y. ATR-FTIR studies in pore forming and membrane induced fusion peptides. Biochim. Biophys. Acta 1828, 2306–2313 (2013)

[77] Subrini O, Sotomayor-Pérez AC, Hessel A, Spiaczka-Karst J, Selwa E, Sapay N, Veneziano R, Pansieri J, Chopineau J, Ladant D, Chenal A. Characterization of a membrane-active peptide from the Bordetella pertussis CyaA toxin. J Biol Chem. 288, 32585-32598 (2013)

[78] Bumba L, Masin J, Fiser R, Sebo P. Bordetella adenylate cyclase toxin mobilizes its beta2 integrin receptor into lipid rafts to accomplish translocation across target cell membrane in two steps. PLoS Pathog. 6, e1000901 (2010)

[79] von Heijne G. Membrane proteins: the amino acid composition of membrane-penetrating segments. Eur J Biochem. 120, 275-278 (1981)

[80] Hildebrand PW, Preissner R, Frömmel C. Structural features of transmembrane helices. FEBS Lett. 559, 145-151 (2004)

[81] Strandberg E, Ozdirekcan S, Rijkers DT, van der Wel PC, Koeppe RE 2nd, Liskamp RM, Killian JA. Tilt angles of transmembrane model peptides in oriented and non-oriented lipid bilayers as determined by 2H solid-state NMR. Biophys J. 86, 3709-3721 (2004)

[82] Duque, D.; Li, X. J.; Katsov, K.; Schick, M. J. Molecular theory of hydrophobic mismatch between lipids and peptides. Chem. Phys. 116, 10478-10484 (2002)

[83] W.-C.C. Hung, M.-T.T. Lee, F.-Y.Y. Chen, H.W. Huang. The condensing effect of cholesterol in lipid bilayers. Biophys. J. 92, 3960-3967 (2007)

[84] Park SH, Opella SJ. Tilt angle of a trans-membrane helix is determined by hydrophobic mismatch. J Mol Biol. 350, 310-318 (2005)

[85] J. Fantini, D. Carlus, N. Yahi. The fusogenic tilted peptide (67–78) of alpha-synuclein is a cholesterol binding domain. Biochim. Biophys. Acta 1808, 2343-2351 (2011)

[86] J. Fantini, N. Yahi. Brain lipids in synaptic function and neurological disease. Clues to Innovative Therapeutic Strategies for Brain Disorders. Elsevier Academic Press, San Francisco ISBN: 9780128001110 (2015)

[87] Koufos E, Chang EH, Rasti ES, Krueger E, Brown AC. Use of a Cholesterol Recognition Amino Acid Consensus Peptide to Inhibit Binding of a Bacterial Toxin to Cholesterol. Biochemistry 55, 4787-97 (2016)

[88] Voegele A, Sadi M, O’Brien DP, Gehan P, Raoux-Barbot D, Davi M, Hoos S, Brûlé S, Raynal B, Weber P, Mechaly A, Haouz A, Rodriguez N, Vachette P, Durand D, Brier S, Ladant D, Chenal A. A High-Affinity Calmodulin-Binding Site in the CyaA Toxin Translocation Domain is Essential for Invasion of Eukaryotic Cells. Adv Sci (Weinh) 8, 2003630 (2021)

